# Translation-dependent mRNA localization to *Caenorhabditis* elegans adherens junctions

**DOI:** 10.1101/2021.05.20.444977

**Authors:** Cristina Tocchini, Michèle Rohner, Stephen E. Von Stetina, Susan E. Mango

## Abstract

mRNA localization is an evolutionarily widespread phenomenon that facilitates sub-cellular protein targeting. Extensive work has focused on mRNA targeting through “zip codes” within untranslated regions (UTRs), while much less is known about translation-dependent cues. Here, we examine mRNA localization in *Caenorhabditis elegans* embryonic epithelia. From an smFISH-based survey, we identified mRNAs associated with the cell membrane or cortex, and with apical junctions in a stage- and cell type-specific manner. Mutational analyses for one of these transcripts, *dlg-1*/*discs large*, revealed that it relied on a translation-dependent process and did not require its 5’ or 3’ UTR. We suggest a model in which *dlg-1* transcripts are co-translationally colocalized with the encoded protein: first the translating complex goes to the cell membrane through sequences of the SH3 domain, and then to the apical junction by the L27 and PDZ sequences. In addition, the Hook and GuK sequences contribute to the second step: they are required for mRNA, but not protein, to accumulate at the apical junctions from locations at or near the membrane. These studies identify a translation-based process for mRNA localization within developing epithelia and determine the necessary cis-acting sequences for *dlg-1* mRNA targeting.

**Summary statement:** An smFISH-based survey identified a subset of mRNAs coding for junctional components that localize at or in the proximity of the adherent junction through a translation-dependent mechanism.

## Introduction

mRNA localization is an efficient means to place the associated translation products in the appropriate subcellular location (Ephrussi et al., 1991; Lecuyer et al., 2007; Takizawa & Herskowitz, 1997). Large-scale studies in diverse organisms have revealed that many mRNAs are enriched at specific subcellular loci (Jambor et al., 2015; Lecuyer et al., 2007). This mechanism is essential to establish embryonic patterning (Frigerio et al., 1986; Rebagliati et al., 1985), distribute determinants asymmetrically in precursor cells (Broadus et al., 1998; Li et al., 1997), and segregate functionally distinct compartments in differentiated and polarized cells like neurons or epithelial cells (Ryder & Lerit, 2018). For example, a global analysis of localized mRNAs in murine intestinal epithelia found that 30% of highly expressed transcripts were polarized (Moor, 2017). The frequent close apposition of mRNAs and their translated proteins indicates that one function of mRNA localization is to enrich proteins at their final destinations through localized translation (Ryder & Lerit, 2018). However, other functions exist, such as targeted protein degradation (Chouaib et al., 2020), translational repression (Kourtidis et al., 2017), and RNA stabilization or storage (Standart & Weil, 2018).

Cells use a variety of mechanisms to position mRNAs within cells. UTRs often harbor localization elements (“zip codes”) that dictate where an mRNA should be delivered (Katz et al., 2012; Kislauskis et al., 1994; Nagaoka et al., 2012). Correct splicing and the presence of exon junction complexes can also play a role in mRNA enrichment to certain cellular subcellular localizations (Hachet & Ephrussi, 2004; Kwon et al., 2021). On the other hand, mRNAs coding for transmembrane or secreted proteins can be localized through a translation-dependent mechanism. For example, the localization factor Signal Recognition Particle (SRP) binds the nascent signal peptide of endoplasmic reticulum (ER)-bound proteins, arrests cytoplasmic translation, and docks at the ER. Translation is resumed after docking, and transmembrane machinery allows the translocation of the fully synthetized proteins into the ER (Weis et al., 2013). More recently, studies with translation inhibitors puromycin and cycloheximide have implicated nascent peptides for mRNA localization to other membranous organelles or non-P body foci, but the exact mechanisms for delivery is unknown for most factors (Chouaib et al., 2020).

In *C. elegans*, mRNA localization has been studied mainly in the context of post-transcriptional gene silencing in membraneless organelles, specifically germline P granules, somatic P-bodies and stress granules, where transcripts are stabilized (Scheckel et al., 2012), protected from degradation or small RNA-mediated gene silencing (Gallo et al., 2008; Ouyang et al., 2019; Shukla et al., 2020), or repressed translationally (Voronina, 2013). In these instances, mRNAs are commonly post-transcriptionally regulated and localized through their 3’UTRs (Parker et al., 2020; Wright et al., 2011). 3’UTR-dependent mRNA localization also occurs in axons of adult neurons (Yan et al., 2009), where mRNA localization is paired with local translation. However, not all localized RNAs rely on their 3’UTRs. A recent study on the early *C. elegans* embryo demonstrated the dispensability of 3’UTR to localize their mRNAs (Parker et al., 2020). However, the mechanisms that localize these RNAs are currently unknown.

In this study, we focused on mRNA localization during development of *C. elegans* embryonic epithelia. *C. elegans* embryogenesis is highly stereotyped, giving rise to an invariant number and positioning of epithelial cells. Epithelial morphogenesis starts from the embryonic stage possessing eight endodermal cells (8E stage) (Sulston et al., 1983) when cell junctions, commonly referred to as the *C. elegans* Apical Junction (*Ce*AJ) (McMahon et al., 2001), begin to form. *Ce*AJs are fully established during the so-called bean and comma stages (names attributed to the early elongation stages based on the shape of the embryo (Sulston et al., 1983). *C. elegans* possesses a single type of apical junction that comprises two adjacent adhesion systems, AS-I and AS-II (Bossinger et al., 2015). AS-I is composed of a Cadherin-Catenin Complex (CCC), constituted by HMR-1/E-Cadherin, HMP-1/α-Catenin, HMP-2/β-Catenin, and JAC-1/p120-Catenin, which link to intermediate filaments of the cytoskeleton and F-actin (Costa et al., 1998; Pettitt et al., 2003). Additional cytoskeletal organizers (e.g., SMA-1) contribute to the correct architecture of AS-I (McKeown et al., 1998). In AS-II, a DLG-1/Discs Large and AJM-1 Complex (DAC) provides a link between the proposed adhesion molecule of the AS-II, called SAX-7/L1CAM (Chen & Zhou, 2010), and cytoskeletal-associated components SMA-1/βH-Spectrin, ERM-1/Ezrin/Radixin/Moesin, and actin filaments (Bernadskaya et al., 2011; Gobel et al., 2004; McKeown et al., 1998; Van Furden et al., 2004). A series of evolutionarily conserved ancillary proteins (actin filaments, claudins, spectrins, PAR proteins, *etc*.) help form and maintain the *Ce*AJs (Armenti & Nance, 2012).

In this work, we conducted a single molecule fluorescent *in situ* hybridization (smFISH)-based survey on the *C. elegans* embryo and tested the localization of mRNAs coding for factors belonging to AS-I and AS-II, as well as for proteins involved in *Ce*AJ formation and maintenance. We identified transcripts enriched at the *Ce*AJ in a stage- and cell type-specific. Genetic and imaging analyses of transgenic lines for one of the identified localized mRNAs, *dlg-1/discs large*, mapped domains required for localization. Our data demonstrate that the *dlg-1* UTRs are dispensable, whereas translation is required for localization.

## Results

### mRNAs coding for the main components of the cell adhesion system II are enriched at the *Ce*AJ

We began our analysis of mRNA localization by surveying twenty-five transcripts that code for the major factors involved in cell polarity and *Ce*AJ formation, as well as some cytoskeletal components (Fig. 1A and Table 1). The protein products of the tested mRNAs are localized differentially along the cell membrane/cortex and cytoplasm (Table 1). We identified epithelial cells and the *Ce*AJ using a CRISPR-engineered DLG-1::GFP fusion (Heppert et al., 2018). Our survey revealed mRNAs with varying degrees of localization within epithelia (Fig. 1B-G). Five of these transcripts were enriched at specific loci at or near the cell membrane: laterally and at the *Ce*AJ for *dlg-1* (Fig. 1C and S1A), solely at the *Ce*AJ for *ajm-1* and *erm-1* (Fig. 1D-E), apically and at the *Ce*AJ for *sma-1* (Fig. 1F), and apically for *vab-10a* (Fig. 1G). Interestingly, all these transcripts but *vab-10a* encode the main cytoplasmic components of the AS-II.

**Figure 1.**
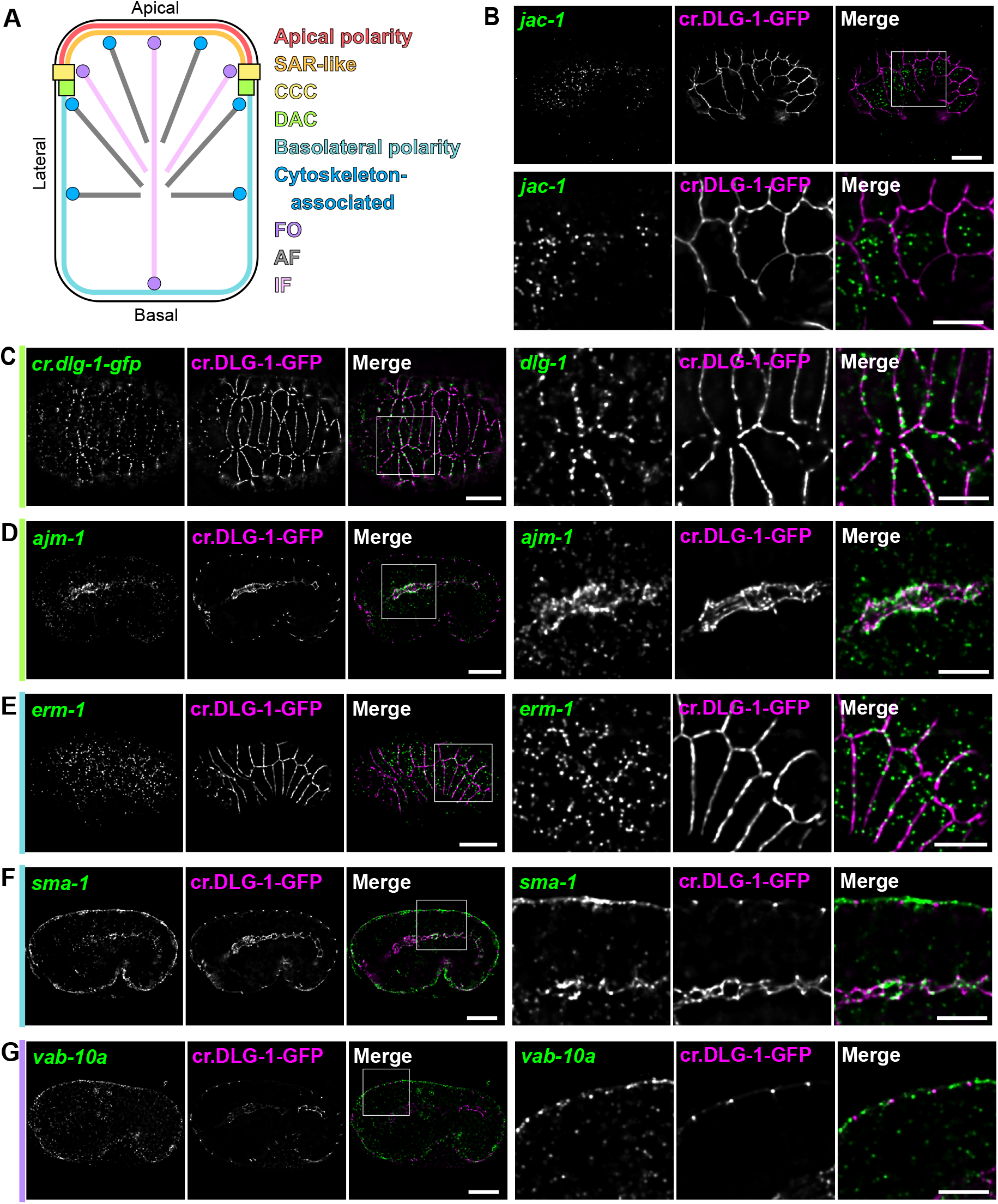
Five mRNAs coding for DAC components, basolateral polarity factors, and a fibrous organelle-bound protein are enriched at the cell membrane. **A**. Simplified color-coded schematics of a *C. elegans* epithelial cell, highlighting the classes of factors involved in apicobasal polarity and maintenance of epithelial morphology. Red: apical polarity factors. Orange: subapical region-like (SAR-like). Yellow: cadherin-catenin complex (CCC). Green: DLG-1/AJM-1 complex (DAC). Cyan: basolateral polarity factors. Blue: cytoskeletal-associated components. Purple: fibrous organelles (FO). Gray: actin filaments (AF). Pink: intermediate filaments (IF). **B**. Fluorescent micrographs of a *C. elegans* embryo at the comma stage (upper panels) and zoom-ins (lower panels) showing smFISH signal of an unlocalized mRNAs (*jac-1* – green), fluorescent signal of the endogenous GFP-tagged DLG-1 protein (cr.DLG-1-GFP, magenta), and merges. **C-G**. Fluorescent micrographs of entire *C. elegans* embryos (left panels) and zoom-ins (right panels) showing smFISH signal of localized mRNAs (*dlg-1* in epidermal cells of a bean stage (C), *ajm-1* in pharyngeal cells of a late comma stage (D), *erm-1* in epidermal cells of a bean stage (E), *sma-1* in epidermal and pharyngeal cells of a late comma stage (F), and *vab-10a* in epidermal cells of a comma stage (G) – green), fluorescent signal of the endogenous GFP-tagged DLG-1 protein (cr.DLG-1-GFP, magenta), and merges. To the left of each image, bars color-coded as in (A) to indicate the sub-class of the factors the mRNAs code for. Squares: portion of the embryo shown in the zoom-ins. Scale bar (entire embryos): 10 µm. Scale bars (zoom-ins): 5 µm.

**Table 1.**
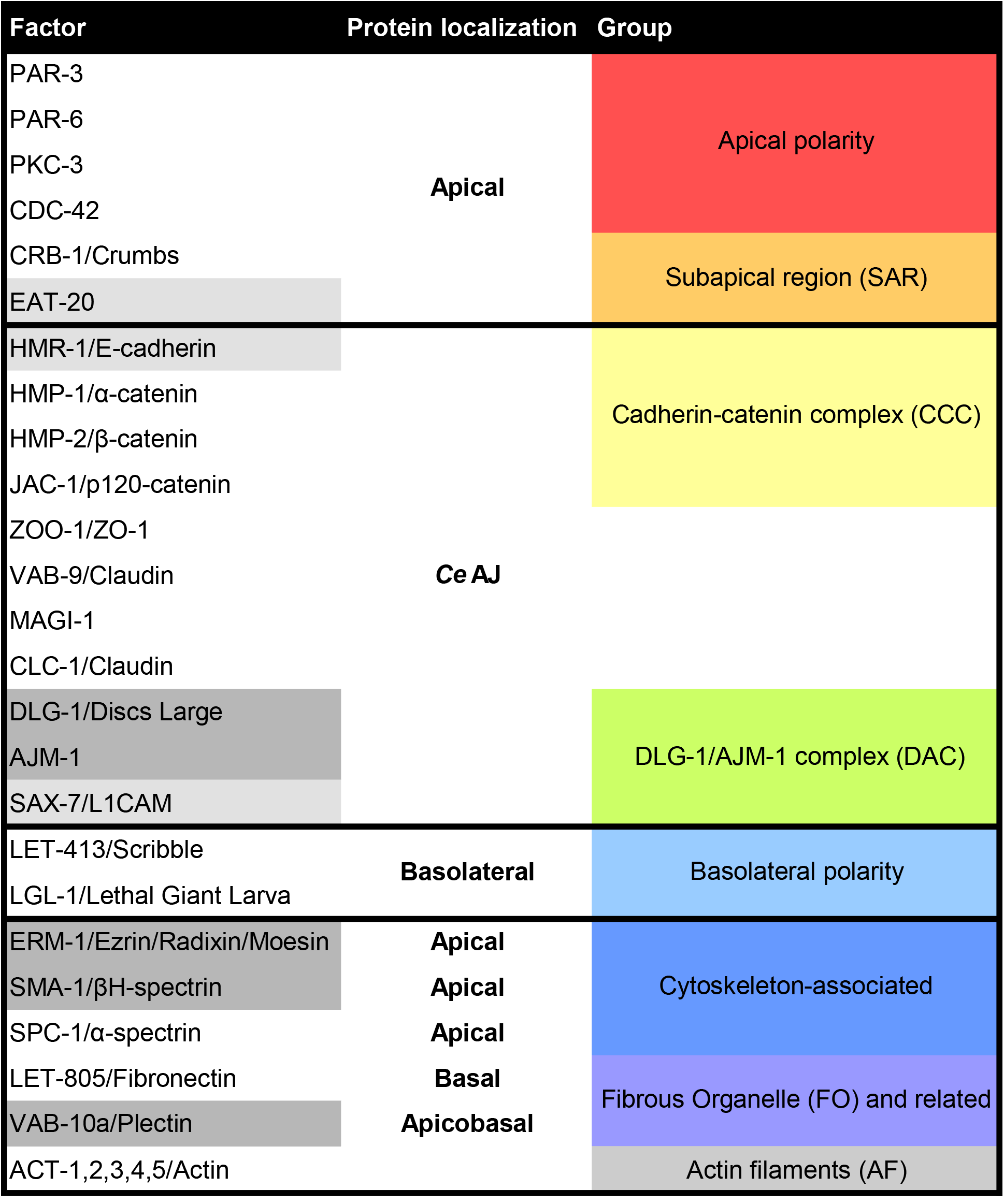
List of factors enrolled in the smFISH survey. Names of the factors and their orthologues whose mRNAs were tested in our smFISH-based survey for localized mRNAs. Additional columns state their subcellular localization and to which group of localized protein they belong to (same color-code as in Fig. 1A). Light gray: perinuclearly localized mRNAs. Dark gray: membrane localized mRNAs.

Beyond AS-II-coding transcripts, our survey also detected three additional mRNAs (*hmr-1, sax-7*, and *eat-20*) that showed some instances of perinuclear localization (Fig. S1B-D). HMR-1/E-cadherin and SAX-7/L1CAM constitute the transmembrane components (putative for SAX-7) of the CCC and the DAC, respectively (Chen & Zhou, 2010; Costa et al., 1998). EAT-20, a Crumbs-like factor involved in apicobasal polarity, is a transmembrane protein as well (Shibata et al., 1999). Bioinformatic analyses of their sequences confirmed the presence of signal peptides in all the three proteins (Methods). Therefore, the perinuclear localization of their transcripts likely reflects classical ER-associated translation (Hermesh & Jansen, 2013).

Taken together, our smFISH survey revealed eight localized mRNAs, five at the cell membrane and three perinuclear. These data indicate that mRNA membrane localization is a feature of the AS-II cell adhesion system, except for the putative transmembrane protein-coding *sax-7* and *actin*.

### *dlg-1* and *ajm-1* mRNA enrichment at the apical junction varies in a stage- and cell type-specific manner

Close examination of the smFISH data showed that mRNA localization varied in a stage- and cell type-specific manner, including transcripts encoding components of the same complex. Specifically, DLG-1 and AJM-1 form a complex (Bossinger et al., 2001) yet differed in the spatiotemporal localization of their mRNAs during epithelial morphogenesis (Fig. 2A). *dlg-1* and *ajm-1* start to be expressed at the 4E embryonic stage. Although epidermal *Ce*AJ (e*Ce*AJ) do not exist yet at the 4E stage (Fig. 2A, upper-most panels), we detected some *dlg-1* mRNA localized near the cell membrane, marked by the basolateral factor LET-413 (Fig. S2A). During e*Ce*AJ formation (8E and 16E stages), when DLG-1 protein is first detectable, *dlg-1* mRNA showed a marked colocalization with its protein, whereas *ajm-1* mRNA was predominantly cytoplasmic (Fig. 2A). When e*Ce*AJ were formed and fully established (bean and comma stages), forming the typical continuous and circumferential belt-like structure at the apical side of the cell membrane, both *dlg-1* and *ajm-1* showed a peak of enrichment at the *Ce*AJ of about 40% (Fig. 2A-B). Afterwards, during later stages of embryonic body elongation (1.5-fold stage), *dlg-1* and *ajm-1* mRNA localization slowly diminished (Fig. 2A-B).

**Figure 2.**
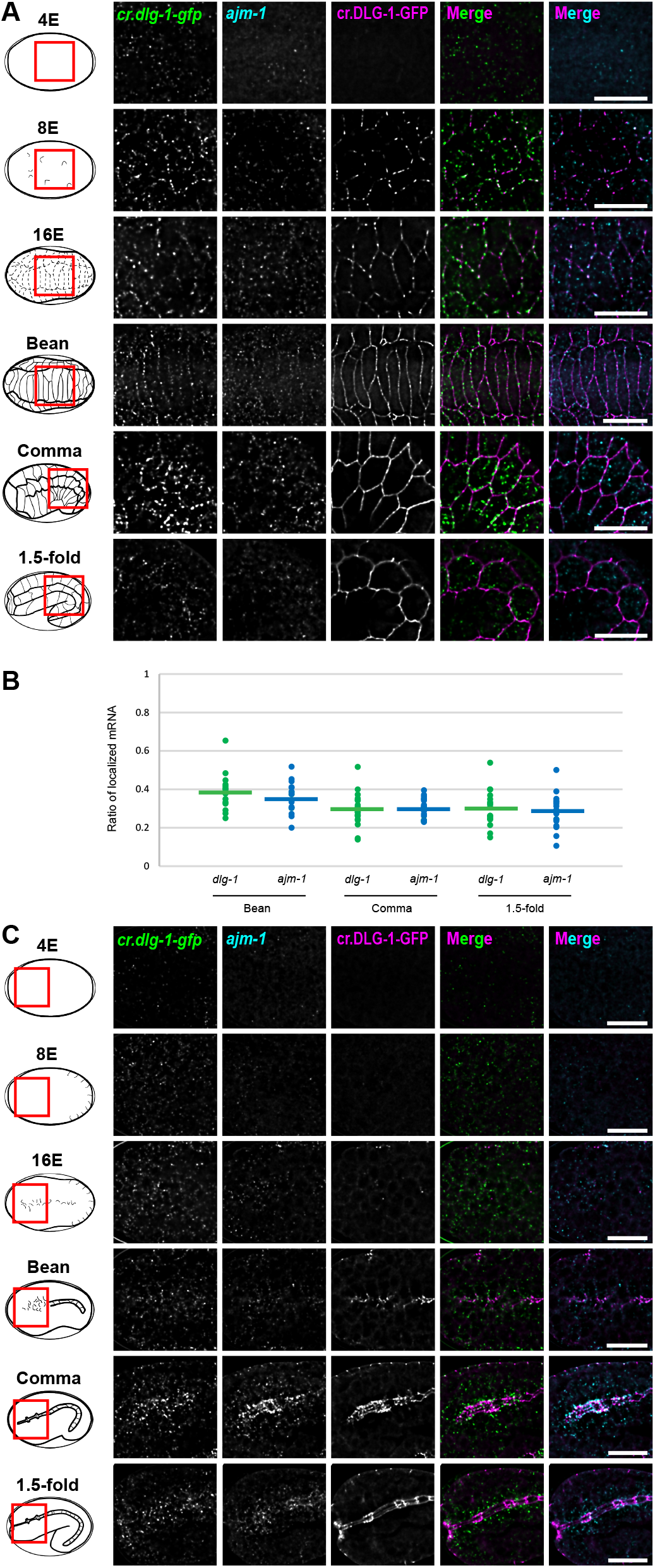
*dlg-1* and *ajm-1* mRNA localization changes dynamically during epithelial morphogenesis. **A**. Left: names and schematics of the analyzed embryonic stages (4E: no junctions; 8E: nascent junctions; 16E: junction maturation; bean: junction formation; comma-1.5-fold: established junction). Red squares: portion of the embryo shown on the right. Right panels: fluorescent micrographs of epidermal and seam cells (A) of *C. elegans* embryos showing smFISH signal of two localized mRNAs, *dlg-1* (*cr*.*dlg-1-gfp*, green) and *ajm-1* (cyan), fluorescent signal of the CRISPR-engineered GFP-tagged DLG-1 protein (cr.DLG-1-GFP, magenta), and merges. Scale bars: 5 µm. **B**. Dot plot: each dot represents the ratio of laterally localized versus total cellular *dlg-1* (green) and *ajm-1* (blue) mRNA (Y axis) in each seam cell analyzed at the stated embryonic stages (X axis: bean (n = 17), comma (n = 20), and 1.5-fold (n = 17)). Horizontal bars represent the mean: *dlg-1* – bean: 0.39 (standard deviation: 0.09); *ajm-1* – bean: 0.36 (standard deviation: 0.08); *dlg-1* – comma: 0.29 (standard deviation (SD): 0.08); *ajm-1* – comma: 0.30 (SD: 0.05); *dlg-1* – 1.5-fold: 0.30 (SD: 0.09); *ajm-1* – 1.5-fold: 0.28 (SD: 0.09). Data derived from 5 different embryos for each stage. **C**. Same as in (A) but in pharyngeal cells.

Analyses of transverse sections of lateral membranes of epidermal (seam) cells at the bean stage demonstrated that *dlg-1* mRNA did not only co-localize with the *Ce*AJ, but it was also present laterally (Fig. S2B). The lateral localization of *dlg-1* mRNA diminished at later stages of development (comma stage) in favor of a more consistent co-localization with the fully mature *Ce*AJ (Fig. S2C).

Morphogenesis of the digestive track showed a different pattern for *dlg-1* and *ajm-1* mRNA junctional localization (Fig. 2B). During foregut or pharyngeal *Ce*AJ (p*Ce*AJ) maturation at the 16E stage, and after full formation at the bean stage, *dlg-1* and *ajm-1* mRNA were only mildly enriched at the membrane (Fig. 2B). Only at the comma stage, when p*Ce*AJ were fully established, a higher degree of localized mRNA could be observed, especially for *ajm-1* mRNA (Fig. 2B). At later stages of pharyngeal morphogenesis (1.5-fold stage), as observed for the epidermis, mRNA enrichment at the p*Ce*AJ decreased gradually (Fig. 2B). These data demonstrate enrichment at the *Ce*AJ for two of our identified localized mRNAs at distinct stages and cell types of embryogenesis.

### The localization of *dlg-1* mRNA at the *Ce*AJ does not depend on its UTRs

mRNA localization commonly involves recognition of zip codes located within UTRs (Chaudhuri et al., 2020; Jambhekar & Derisi, 2007). To test whether the localization of one of the identified localized mRNAs, *dlg-1*, relied on zip codes, we generated transgenic lines carrying a *dlg-1* gene whose open reading frame (ORF) was fused to GFP and to exogenous UTRs. We used UTRs from mRNAs that do not localize near cell membranes, namely *sax-7* and *unc-54* (Fig. S3A-B). One construct (“3’UTR” reporter) substituted the endogenous *dlg-1* 3’UTR with an *unc-54* 3’UTR (Lockwood et al., 2008; McMahon et al., 2001), and a second construct exchanged both the endogenous 5’ and 3’UTRs (“5’-3’UTRs” reporter) by additionally substituting the endogenous *dlg-1* 5’UTR with a *sax-7* 5’UTR to the 3’UTR reporter construct (Fig. 3A). Given that the transgenic constructs were expressed in a wild-type background, smFISH experiments were conducted with probes against the GFP RNA sequence to assess specifically the localization of the transgenic *dlg-1::*GFP mRNAs (*tg*.*dlg-1*). The mRNA localization patterns of the two UTR reporters were compared to the localization of *tg*.*dlg-1* transcripts from a CRISPR line (“wild-type”, Fig. 3A; Heppert et al., 2018). Both reporter strains showed enrichment and localization dynamics of their transcripts that were comparable to the wild-type *tg*.*dlg-1* (Fig. 3B). These results indicate that the UTR sequences of *dlg-1* mRNA are not required for its localization.

**Figure 3.**
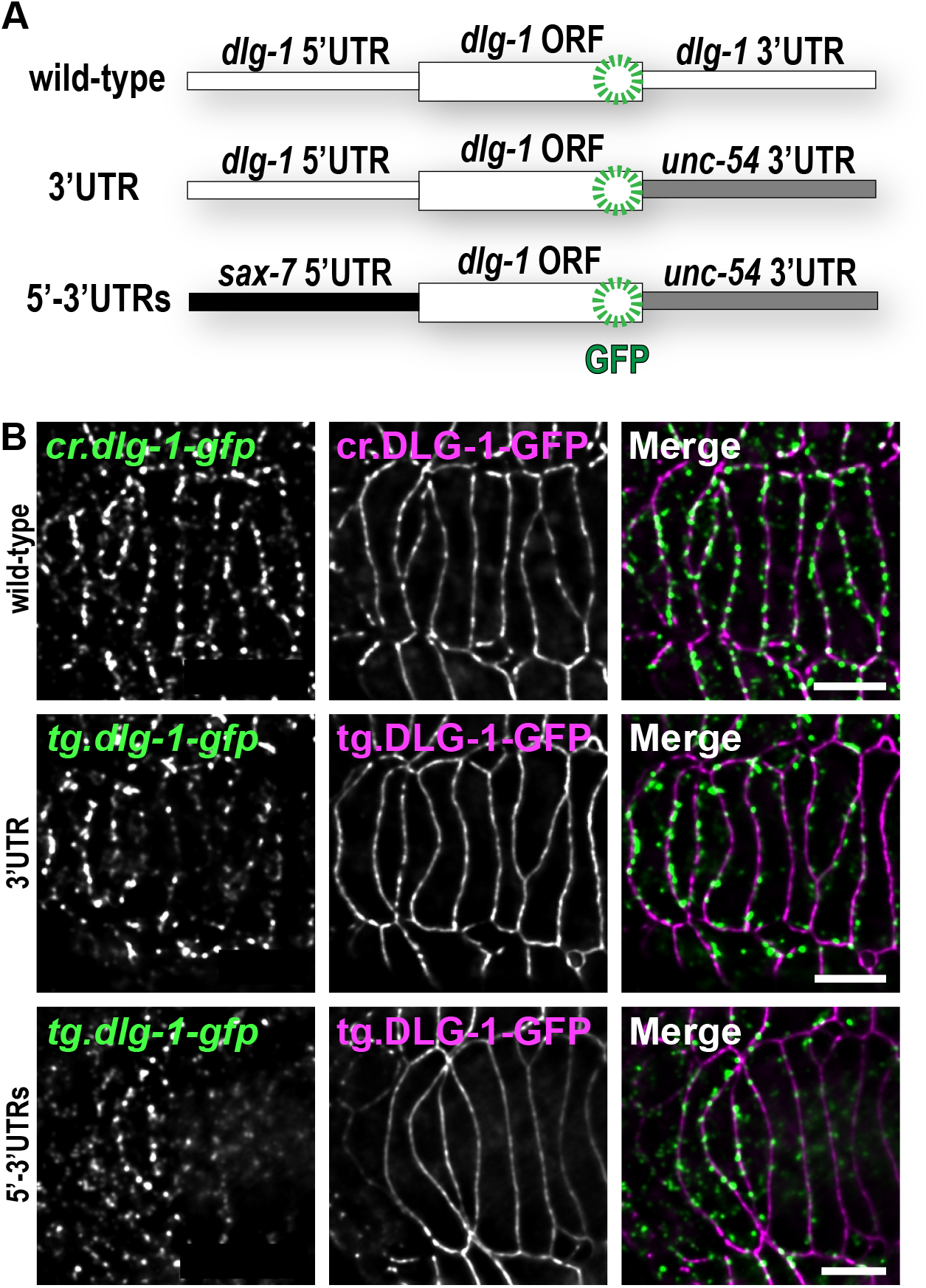
*dlg-1* endogenous 5’ and 3’UTR are not required for its localization. **A**. Schematic representations of the three analyzed transgenes carrying a GFP-tagged *dlg-1* ORF combined with endogenous or exogenous UTRs that are not competent to localize their own mRNAs. “Wild-type”: CRISPR-engineered line with endogenous *dlg-1* 5’ and 3’UTRs (white). “3’UTR”: multicopy extrachromosomal transgenic line with *dlg-1* 5’UTR (white) and *unc-54* 3’UTR (gray). “5’-3’UTRs”: multicopy extrachromosomal transgenic line with *sax-7* 5’UTR (black) and *unc-54* 3’UTR (gray). **B**. Fluorescent micrographs of a dorsal portion of epithelial cells at the bean stage of *C. elegans* embryos showing smFISH signal of CRISPR or transgenic *dlg-1* mRNAs (*cr*.*dlg-1-gfp* and *tg*.*dlg-1-gfp*, green), fluorescent signal of CRISPR or transgenic GFP-tagged DLG-1 protein (cr.DLG-1-GFP and tg.DLG-1-GFP, magenta), and merges. Scale bars: 5 µm.

### The localization of *dlg-1* mRNA at the *Ce*AJ is translation-dependent

Co-translational mechanisms for mRNA delivery have been described for mRNA encoding transmembrane and secreted proteins (Nyathi et al., 2013). Recent studies have suggested that co-translational mRNA localization can also exist for transcripts encoding proteins in other subcellular locations (Chouaib et al., 2020; Hirashima et al., 2018; Safieddine et al., 2021). To determine whether *dlg-1* mRNA localization occurs co-translationally, we designed a transgene to inhibit normal translation by deleting two nucleotides (TG) within the start codon of an otherwise wild-type sequence (Fig. 4A) and generated two transgenic lines. Ribosomes scanning the mRNA from the 5’ end would encounter two new AUG start codons that are each out-of-frame compared to the wild-type (Fig. 4B). The first in-frame AUG after the deletion is located at position 47 (Fig. 4A-B and S4A). We generated transgenic lines in a *smg-2* mutant strain (Hodgkin et al., 1989) to avoid mRNA degradation by nonsense-mediated decay (NMD), which recognizes and destroys mRNAs with precocious translation termination. As a control, we verified by smFISH that wild-type *tg*.*dlg-1* mRNA was localized normally in *smg-2* mutant embryos, demonstrating that NMD does not interfere with targeting *dlg-1* transcripts to the *Ce*AJ (Figure 4C-D).

**Figure 4.**
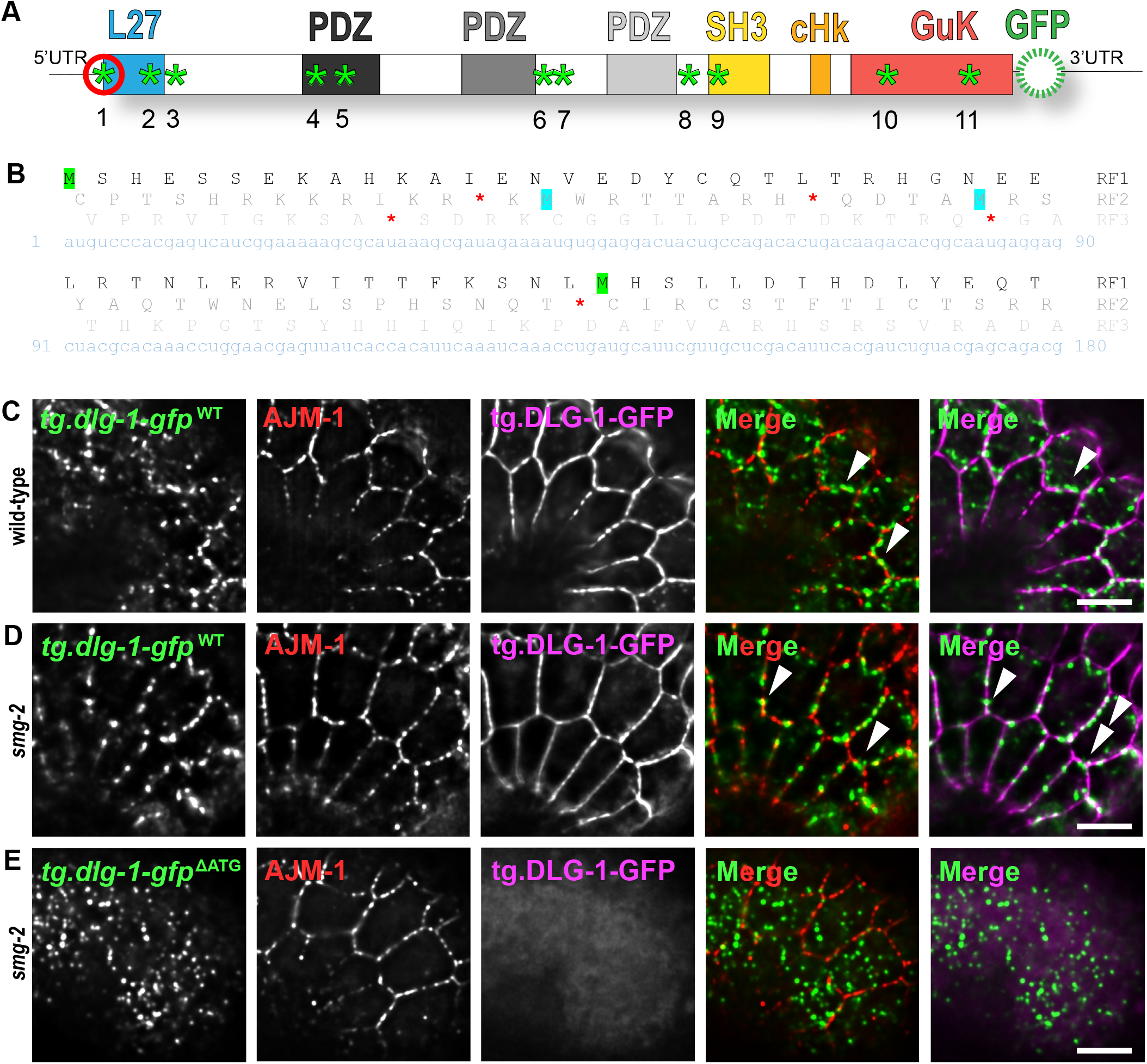
*dlg-1* mRNA localization depend on its translation. **A**. Schematic representation of transgenic *dlg-1-gfp* mRNA. Green asterisks: possible in-frame AUG. Red circle: the first AUG whose corresponding TG nucleotides were elicited from the transgenic “ΔATG” sequence. **B**. Primary sequence of the first 180 nucleotides of *dlg-1* CDS (light blue), and its corresponding translation into its three reading frames (RF1-3; shades of gray). Highlighted in green: the first two methionines of the main frame (RF1). In cyan: the first two methionines in the second frame (RF2). Red asterisks: termination codons in every frame. **C-E**. Fluorescent micrographs of multicopy extrachromosomal transgenic lines of a lateral portion of seam and epidermal cells at a late bean stage of *C. elegans* embryos. smFISH signal of wild-type (“WT” (B-C)) and altered ATG (“ΔATG” (D)) *tg*.*dlg-1-gfp* mRNAs (green), immuno-fluorescent signal of the endogenous AJM-1 protein (red), fluorescent signal of the corresponding tg.DLG-1-GFP protein (magenta), and merges. Corresponding genotypes are on the right. The wild-type transgene in (B) is expressed in wild-type animals. WT and ΔATG transgenes in (C) and (D) are expressed in a null mutant for an NMD component (*smg-2*). Arrowheads indicate examples of localized mRNA. Scale bars: 5 µm.

Next, we examined the localization of our non-translatable *dlg-1::gfp* mRNA (“ΔATG”) in *smg-2* mutant embryos with smFISH paired to fluorescence analysis for DLG-1::GFP protein, to tract the degree of in-frame translation. The C-terminal position of the GFP moiety ensures that mRNAs that initiate in-frame translation anywhere within the DLG-1 coding sequence are GFP-positive. AJM-1 antibody staining was used to identify the *Ce*AJ. The embryos of one of our ΔATG transgenic lines lacked any detectable DLG-1::GFP protein and displayed a marked decrease of mRNA at the *Ce*AJ compared to controls (Fig. 4E). We conclude that translation is required for mRNA localization. Embryos from our second ΔATG transgenic line displayed a little GFP protein and some *dlg-1::gfp* mRNA. We speculate that truncated DLG-1 protein may be generated by one or more of the ten alternative in-frame AUGs that can be found within the *dlg-1* mRNA (Fig. 4A and S4A; Table S1). For the second line, we observed some mRNA localized at the *Ce*AJ, and it was always in proximity to DLG-1::GFP protein (Fig. S4B). These data suggest that *dlg-1* mRNA localization depends on its ongoing translation (*e*.*g*., line 1; Fig. 4E), and that even low amounts of translational activity are sufficient for mRNA delivery to its final location (e.g., line 2; Fig. S4B).

### The C-terminus is necessary and sufficient for mRNA localization to the membrane, and the N-terminus is important for enrichment at the junction

Our data suggested that translation of *dlg-1* mRNA in cis lead to its enrichment at the *Ce*AJ. DLG-1 is a complex protein with different domains that allow its localization and function, as diagrammed in figure 5 (Fig. 5A; (Firestein & Rongo, 2001). To define critical regions for mRNA localization, we deleted these domains using existing (Firestein & Rongo, 2001; Lockwood et al., 2008) and newly generated transgenic lines (Methods) (Table 2). Immuno-staining for endogenous AJM-1 provided a spatial reference for the localization of the *Ce*AJ that was unaffected in any of our analyzed transgenic strains, which also expressed endogenous, wild-type DLG-1. smFISH with GFP probes were specific for transgenic *dlg-1*::GFP (“*tg*.*dlg-1*”), and GFP fluorescence from transgenic DLG-1 (“tg.DLG-1”) provided a readout for the localization of the transgenic protein. We analyzed images of frontal plane views of epidermal seam cells at the bean stage, paired to apicobasal and apical intensity profile analyses (Fig. 5B-C). Fluorescent images of our full-length control showed lateral and *Ce*AJ enrichment of *tg*.*dlg-1*^*FL*^ mRNA and *Ce*AJ localization for the corresponding tg.DLG-1^FL^ protein (Fig. 5D). These data were confirmed by intensity profile analyses where mRNA peaks (green) largely overlapped with protein peaks (magenta), and these were located at the *Ce*AJ (pink vertical lines) in both apicobasal and apical profiles (Fig. 5D’-’’). A few minor mRNA peaks were observed in the apicobasal intensity profile in between the *Ce*AJ peaks (*i*.*e*., cytoplasmic mRNAs; Fig. 5D’), as well as in the close proximity to the *Ce*AJ peaks in the apical intensity profile (Fig. 5D’’). Such a lateral and *Ce*AJ enrichment of *tg*.*dlg-1*^*FL*^ mRNA resembles what we observed with the CRISPR line (Fig. S2B). The same lateral and junctional transgenic mRNA enrichment and protein localization could also be observed from top views of seam cells in embryos at the same stage (Fig. S5A-A’).

**Table 2.** List of transgenic lines with truncations within the *dlg-1* CDS. Strain names, domains contained, specific amino acid locations, full genotypes, and references for the transgenic *dlg-1*::GFP lines used in this study.

| Name | Domains | Amino acids | Genotype | Reference |
| --- | --- | --- | --- | --- |
| SM2637 | Full-Length (FL) | DLG-1(1-929) | *pxEx636[dlg-1p::dlg-1 5'UTR::dlg-1::GFP::unc-54 3'UTR; rol-6(su1006)]* | This study |
| SM2684 | ΔL27 | DLG-1(Δ17-116) | *jcEx128[dlg-1p::dlg-1(Δ17-116)::GFP::unc-54 3'UTR; rol-6(su1006)]* | Lockwood et al., 2008 |
| SM2641 | ΔL27-PDZs | DLG-1(593-929) | *pxEx640[dlg-1p::dlg-1(593-929)::GFP::unc-54 3'UTR; rol-6(su1006)]* | This study |
| SU359 | ΔHk-GuK | DLG-1(1-710) | *dlg-1(ok318)/dpy-8(q287) X; jcEx94[dlg-1p::dlg-1(1-710)::GFP::unc-54 3'UTR; rol-6(su1006)]* | Lockwood et al., 2008 |
| SM2685 | L27-PDZ1 | DLG-1(1-302) | *jcEx95[dlg-1p::dlg-1(1-302)::GFP::unc-54 3'UTR; rol-6(su1006)]* | Lockwood et al., 2008 |
| SU361 | L27-PDZ1/2 | DLG-1(1-468) | *dlg-1(ok318)/dpy-8(q287) X; jcEx96[dlg-1p::dlg-1(1-468)::GFP::unc-54 3'UTR; rol-6(su1006)]* | Lockwood et al., 2008 |

**Figure 5.**
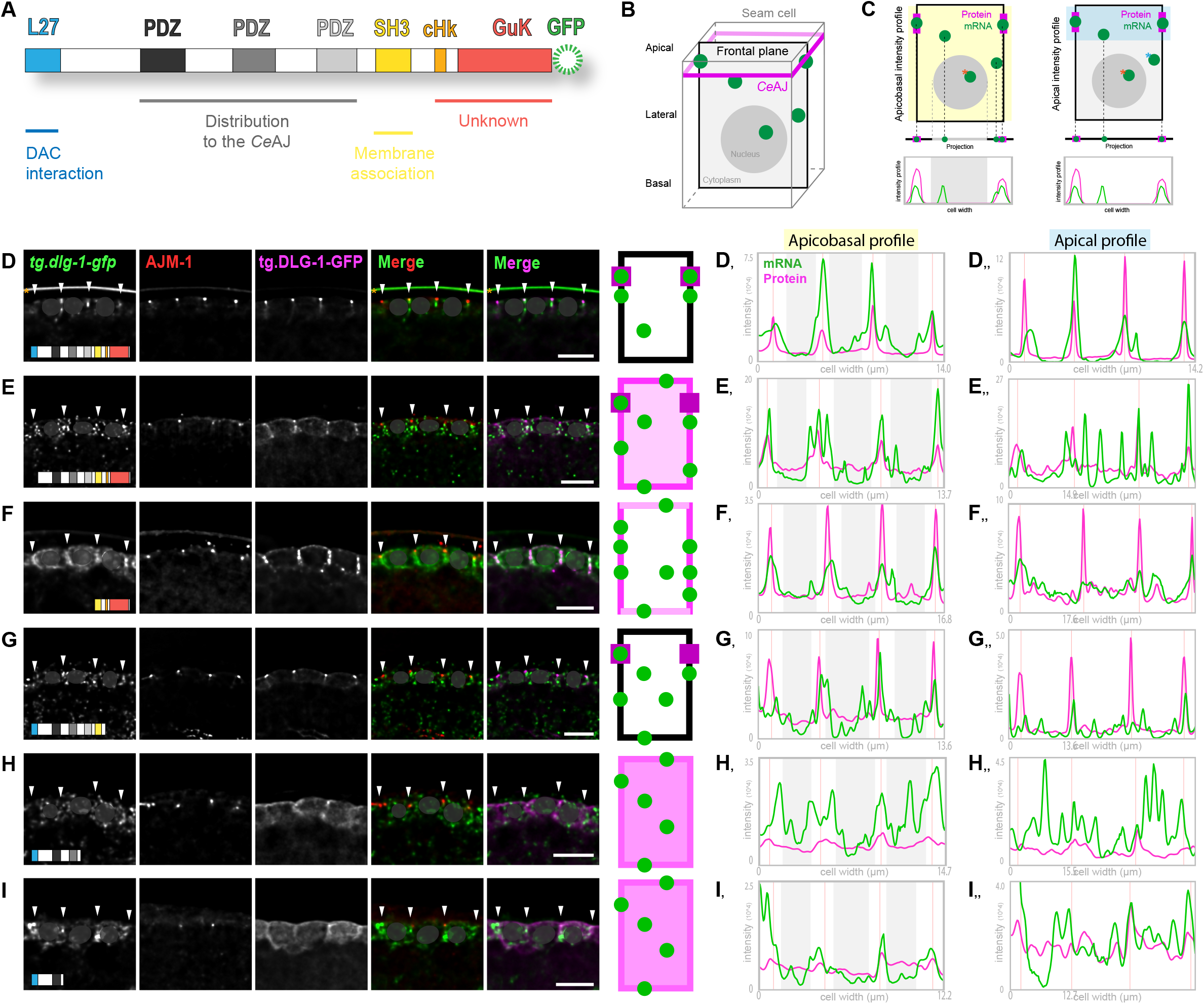
Specific domain-coding sequences of *dlg-1* mRNA are required for its normal localization. **A**. Schematic representation of the full-length transgenic DLG-1 protein, highlighting domains and their known functions (Firestein et al., 2001). Blue: L27 domain. Shades of gray: the three PDZ domains. Yellow: SH3 domain. Orange: the conserved stretch of the Hook domain (cHk). Red: GuK domain. Green circle: GFP, C-terminally tagged. **B**. Schematic representation of a seam cell in 3D (gray cube). Magenta apical belt: *Ce*AJ. A black rectangle shows a frontal plane view in the middle of the cell used to analyze the images in the rest of the figure. Light gray represents the cytoplasm, a dark gray filled circle the nucleus, and green filled circles mRNAs. **C**. Top: simplified schematics of the frontal view of a seam cell (B). Green circles: transgenic *dlg-1* mRNA. Magenta rectangles: transgenic DLG-1 protein. Highlighted in yellow (top left) and in blue (top right) the regions of the cell used for apicobasal and apical intensity profile analyses, respectively. Orange asterisks: mRNAs in the nuclei that have not been considered in the intensity profile analyses, as representing transcription sites. Blue asterisk: example of a cytoplasmic mRNA that would be considered in the apicobasal analysis, but not in the apical. Below: projections of mRNA and protein (and nucleus in the left side) present in the schematics above (same color-code). Bottom: exemplified intensity profile graphs based on the projections above, where peaks show the positions of transgenic *dlg-1* mRNA (green line) and transgenic DLG-1 protein (magenta). X axis: width of the cell; Y axis: fluorescent intensity. The gray box in the left graph represents the projection of the nucleus whose intensities have been removed from the analysis. **D-I**. Frontal plane views of fluorescent micrographs of three adjacent seam cells at the bean stage of *C. elegans* embryos showing smFISH signal of transgenic *tg*.*dlg-1-gfp* mRNAs (full length (D), ΔL27 (E), SH3-cHk-GuK (F), ΔcHk-GuK (G), L27-PDZ1/2 (H), and L27-PDZ1 (I) (green)), immuno-fluorescent signal of the endogenous AJM-1 protein (red), fluorescent signal of the corresponding transgenic GFP-tagged DLG-1 protein (magenta), and merges. Arrowheads: *Ce*AJ localization. Shaded gray circles: the nuclear regions that have been excluded from the intensity profile analyses. Orange asterisks: unspecific signal staining the eggshell. Scale bars: 5 µm. Right side: simplified schematics based on the fluorescent images on the right. Frontal view of a seam cell (rectangle) modelling transgenic mRNA and protein localizations. Green circles: transgenic *dlg-1* mRNA. Shades of magenta: varying degrees of transgenic DLG-1 protein along the membrane (borders) and in the cytoplasm (middle part). **D’-I’**. Intensity profile graphs of three contiguous cells (apicobasal profile (highlighted in yellow) explained in (C) (left side) for the corresponding fluorescent images on the left (D-I). X axis: cell width (µm); Y axis: measured fluorescent intensities. Green lines: transgenic *dlg-1* mRNA intensities. Magenta lines: transgenic DLG-1 protein intensities. Pink vertical lines: location of the *Ce*AJ, identified by peak values for the intensity profile of AJM-1 fluorescent signal (not shown). Light gray panels: nuclei locations that have been evicted from the mRNA channel to avoid quantification of transcriptional signal. **D’’-I’’**. Intensity profile graphs of the sole apical part of the same cells analyzed on the left (apical profile (highlighted in blue) explained in (C – right side)). Axes and color-codes as in (D’-I’).

First, we examined the amino terminus, specifically deletion of the sole L27 domain (ΔL27). This domain is involved in DLG-1 multimerization as well as interactions with the other DAC component, AJM-1 (Firestein & Rongo, 2001; Lockwood et al., 2008). tg.DLG-1^ΔL27^ was localized more broadly along the whole membrane compared to the full-length, although retaining a partial enrichment at the *Ce*AJ (Fig. 5E). Similar to its encoded protein, *tg*.*dlg-1*^*ΔL27*^ mRNA distribution was scattered along the whole membrane (Fig. 5E and Fig. S5B). Intensity profile analyses confirmed the broader membrane distribution of tg.DLG-1^ΔL27^ and the presence of *tg*.*dlg-1*^*ΔL27*^ mRNA along the lateral (and partially apical and basal) membranes. We also observed a small enrichment of *tg*.*dlg-1* mRNA at the *Ce*AJ, less than the wild-type, as well as an increased amount of cytoplasmic mRNA that was greater than the wild-type (Fig. E’-’’). These data suggest that the L27 domain contributes to the accumulation of *dlg-1* mRNA and protein at the *Ce*AJ.

A larger amino-terminal truncation removed the PDZ domains as well as the L27 domain, but left the SH3, Hook, and GuK domains. The SH3 domain is involved in interactions with lateral factors (Lockwood et al., 2008), whereas the Hook and GuK domains have no known role in protein localization (Lockwood et al., 2008). This truncated tg.DLG-1^ΔL27-PDZs^ protein was broadly distributed at the lateral membrane without any enrichment at the *Ce*AJ. *tg*.*dlg-1*1^*ΔL27-PDZs*^ mRNA was distributed all along the membrane, especially laterally (Fig. 5F, 5F’, and S5C). These data show that the carboxy-terminal sequences are sufficient to direct *dlg-1* RNA and protein to the membrane, but that the amino-terminal L27 and PDZ domains are important for targeting the *Ce*AJ (Fig. 5F and 5F’’).

### The Hook and GuK domains uncouple mRNA and protein localization

mRNA and protein localization correlate for the constructs described above, suggesting that either protein dictates mRNA localization or translation occurs at defined locations. This correlation was lost for constructs lacking the Hook and GuK domain (Fig. 5G and 5G’-’’). Deletion of a large part of the Hook domain and the whole GuK domain didn’t affect tg.DLG-1^ΔHk-GuK^ protein localization but did impair *Ce*AJ enrichment of the mRNA (Fig. 5G and 5G’-’’). The mRNA was detected at all membrane surfaces: lateral but also apical and basal (Fig. 5G and S5D). Other mRNA was detected adjacent to the membrane, but not overlapping, and a small proportion was cytoplasmic (Fig. 5G’ and S5D) However, the truncation impaired *Ce*AJ enrichment of *tg*.*dlg-1*^*ΔHk-GuK*^ mRNA (Fig. 5G and 5G’’) and partially affected its lateral localization These data suggest that sequences within the Hook and GuK domains help target *dlg-1* mRNA to the *Ce*AJ from membrane locations and, at least in part, it may also contribute to targeting to the membrane (see Discussion).

Further removal of sequences from the carboxy-terminus impaired tg.DLG-1 protein and *tg*.*dlg-1* mRNA localization. We examined constructs terminated at the second or at the third PDZ domain. Both deletions produced transgenic protein and mRNA present all over the membrane and in the cytoplasm (Fig. 5H-I and S5E). In intensity profile analysis, mRNA and protein peaks showed minimal overlap, confirming the broader and more randomized distribution observed in the fluorescent images (Fig. 5H’-’’, 5I’-’’). These data suggest that *dlg-1* mRNA and protein can accumulate in ectopic locations without degradation, at least the amino terminal half of the protein.

In conclusion, the structure-function analyses revealed that the Hook and GuK domains were required to localize *dlg-1* mRNA, but not protein. The Hook and GuK domains, together with the SH3 domain, were sufficient for both protein and mRNA localization to the lateral membranes. In addition, PDZ domains together with the L27 were necessary, but not sufficient, to bring DLG-1 and *dlg-1* to the *Ce*AJ (Fig. 6).

**Figure 6.**
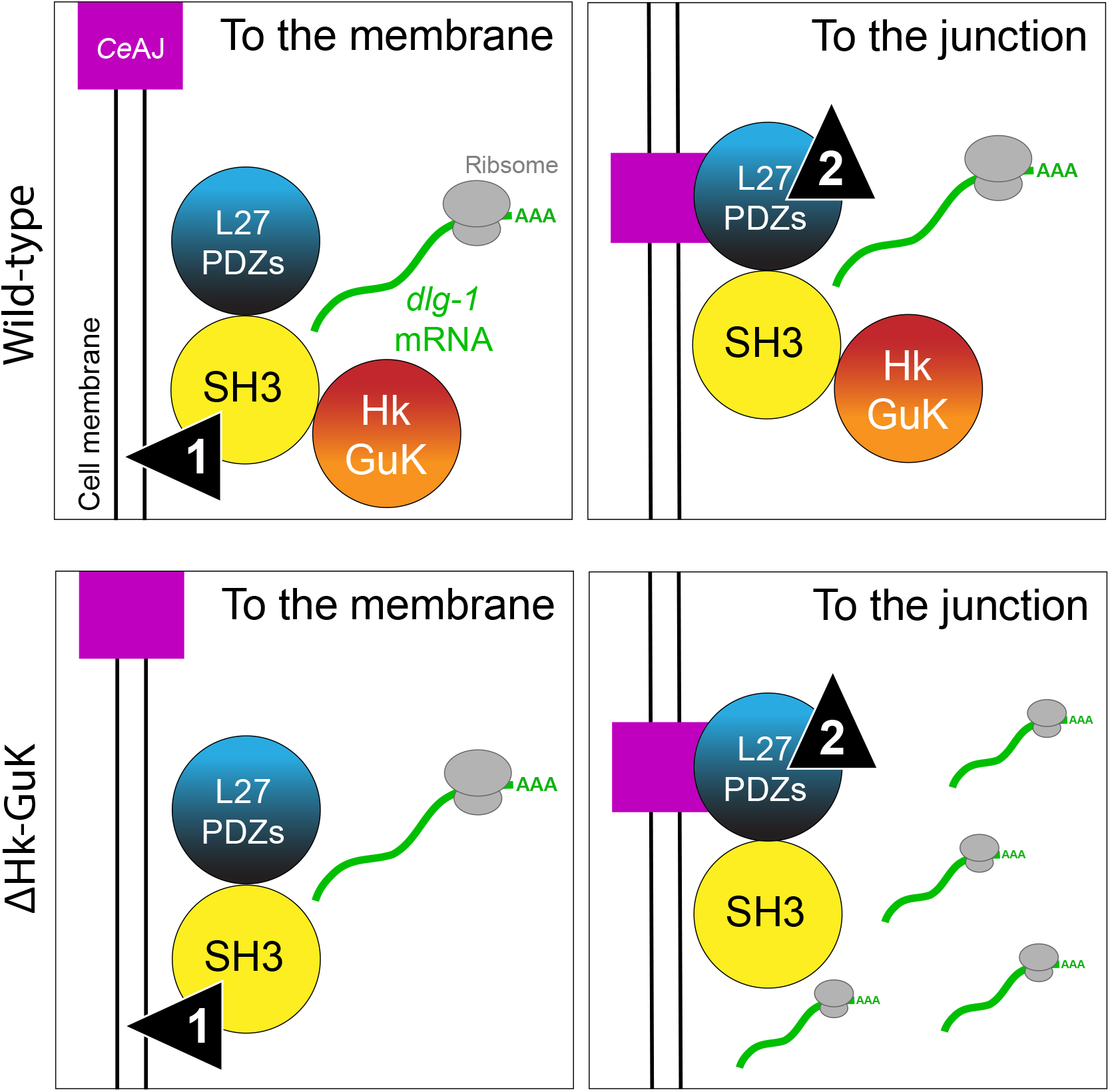
Model for *dlg-1* mRNA localization dependent on Hook and GuK coding sequences. A model showing schematic representations of wild-type (upper panels) and a *dlg-1* truncated version (lower panels) lacking sequences coding for Hook and GuK domains (ΔHk-GuK). Colored circles represent functional sub-units of the DLG-1 protein with the same color-code shown in Fig. 5A: SH3 allows the localization “to the membrane”, and L27 and PDZ domains “to the junction”. Apical junctions are represented as a magenta square at the schematic cell membrane. A translating complex is represented by *dlg-1* mRNA (green) with a ribosome (gray) on top and localized next to the schematic protein (upper panels). Such a translating complex is absent in ΔHk-GuK, where *dlg-1* mRNA is dispersed.

## Discussion

This study has made three contributions towards understanding RNA localization in *C. elegans* embryos. First, we identified mRNAs within adhesion system II of the *Ce*AJ that are localized within epithelial cells. Second, localization of *dlg-1* depends on translation in cis, and not UTR zip codes. Third, specific regions within *dlg-1*/DLG-1 dictate localization at the membrane and, separately, to the *Ce*AJ.

We conducted an smFISH-based survey, which identified endogenous mRNAs that are localized at or near the cell membrane (five out of twenty-five tested mRNAs). mRNA localization in *C. elegans* has only been addressed in the early embryo for maternally provided mRNAs (Parker et al., 2020) and synapses in adult neurons (Yan et al., 2009). Previous studies of *Drosophila* embryogenesis have shown that many mRNAs are localized subcellularly, including mRNAs coding for cytoskeletal and junctional components (Jambor et al., 2015; Lecuyer et al., 2007; Ryder & Lerit, 2018). Among them, *β-actin, E-cadherin*, and *zo-1* mRNAs are localized at the cell cortex of epithelial cells (Ryder & Lerit, 2018). We did not observe orthologues of these mRNAs being cortically localized, suggesting species or cell-type specific differences. Nevertheless, we identified other localized mRNAs, four of which (*dlg-1, ajm-1, erm-1*, and *sma-1*) code for factors that are functionally linked within the AS-II. mRNAs coded by orthologues of *dlg-1* also show a defined subcellular localization in other species in polarized cells such as embryonic cells or neurons. For example, *Drosophila dlg1* mRNA associates transiently to membranes during embryogenesis (mitotic cycle 14): laterally at stage 5 (cellularization) when membranes start to form around nuclei, all around the cell membrane at stages 6 and 7 (cellularization and gastrulation), and it becomes unlocalized at later stages (Lecuyer et al., 2007). In zebrafish, instead, neuronal *dlg1* mRNA localizes stably in myelin sheets of fully differentiated oligodendrocytes (Yergert et al., 2021). These data suggest that *dlg1* mRNA localization may have pivotal roles in development. While *dlg-1* is the most broadly studied, one other mRNA uncovered in our survey, *erm-1*, has been shown to localize in another context. Specifically, *erm-1* mRNA is targeted to the cell periphery in the early *C. elegans* embryo (Parker et al., 2020), providing an additional example of membrane enriched mRNA in different developmental contexts.

Localization of *dlg-1* mRNA depended on its translation rather than its UTRs. Substitution of both the 5’ and 3’ UTRs with UTRs of unlocalized mRNAs did not disrupt association of *dlg-1* mRNA with the *Ce*AJ. This result mirrors orthologues of *dlg-1* in other species, which also do not require their UTRs for subcellular localization (Yergert et al., 2021). Thus, *dlg1* mRNAs are frequently localized within cells, but the mechanism of UTR-independent targeting is unknown in any species. We found that *dlg-1* mRNA required its coding sequences and translation. We analyzed two *C. elegans* lines expressing a full-length *dlg-1*::GFP with the normal ATG deleted (ΔATG). One of these produced no protein and had no mRNA localization, demonstrating the importance of translation in *cis*. Moreover, this line demonstrated that DLG-1 protein supplied in trans was not sufficient for targeting to the *Ce*AJ, as all our transgenic lines were analyzed in embryos expressing wild-type, endogenous DLG-1. The second line fortuitously produced a little protein in some cells. In these expressing cells, both protein and mRNA were localized, suggesting that even a little translation was sufficient to target *dlg-1* mRNA to membrane and *Ce*AJ.

Besides the ATG mutation, the remainder of the *dlg-1*^*ΔATG*^ gene was wild-type, including the intron-exon sequences. This wild-type configuration reveals that sequences and complexes associated with the exon junctions (EJC) are not sufficient to localize *dlg-1* mRNA. Thus, *dlg-1* likely differs from mRNAs like *oskar* in *Drosophila*, where splicing generates a localization element and EJC binding site, which together target *oskar* mRNA within oocytes (Ghosh et al., 2012).

Our structure function analyses highlighted two pathways for mRNA localization: one dependent on the L27-PDZ and SH3 domains and one dependent on the Hook and GuK domains. The first pathway depended on protein sequences known to target DLG-1 protein to the *Ce*AJ. Previous studies had shown that L27 and PDZ domains are key elements for localizing DLG-1 protein as they allow interaction with DAC components and mediate its association with the *Ce*AJ, respectively (Lockwood et al., 2008). In addition, the SH3 domain is required for lateral distribution of DLG-1 protein (Lockwood et al., 2008) which is subsequently redirected to the *Ce*AJ by LET-413 (McMahon et al., 2001). The involvement of these sequences on protein localization and the possible tethering of *dlg-1* mRNA to protein could explain the reliance of *dlg-1* mRNA localization on translation.

We uncovered a second localization pathway for *dlg-1* mRNA that depends on the Hook and GuK sequences. Removal of Hook and GuK sequences mislocalized *dlg-1* mRNA to membranes and occasionally the cytoplasm, without disturbing DLG-1 protein at the *Ce*AJ. Despite normal protein localization, previous studies showed that the Hook and GuK are essential for viability, underscoring their importance(Lockwood et al., 2008). The ability of DLG-1 protein to localize to the *Ce*AJ in the absence of robust mRNA localization may reflect protein targeting by the first pathway. Normally, DLG-1 associates with itself and with AJM-1 via the L27 domain (Lockwood et al., 2008), which could explain the enrichment of mutant DLG-1 from membrane locations to the *Ce*AJ. A second not mutually exclusive possibility is that the small amount of mutant *dlg-1* mRNA at the *Ce*AJ may be translated preferentially.

Our findings suggest a two-step model for targeting DLG-1 to the *Ce*AJ, in which DLG-1 first associates with membranes in an SH3-dependent manner, and a second step where DLG-1 moves to the *Ce*AJ in a process that requires the L27 and, possibly, one or more PDZs. The Hook and GuK domains appear to act predominantly in the second step, to focus *dlg-1* mRNA at the *Ce*AJ. None of these domain combinations was sufficient to target *dlg-1* mRNA to the junction, suggesting these processes synergize to place *dlg-1* mRNA at the *Ce*AJ during embryonic epithelium formation.

## Materials and methods

### Nematode culture

All animal strains were maintained as previously described (Brenner, 1974) at 20°C. Transgenic lines containing extrachromosomal arrays were grown at 15°C to reduce transgene over-expression. Some lines containing extrachromosomal arrays presented instances of mosaicism and differential expression levels among cells due to their extrachromosomal nature. Therefore, we focused on cells with consistent patterns of expression. For a full list of alleles and transgenic lines, see Table S2.

### Generation of transgenic lines

*dlg-1* deletion constructs were generated by overlap extension PCR using pML902 as a template. For *dlg-1* TSLAS and 5’UTR replacement construct, *sax-7* TSLAS and 5’UTR fragments were synthesized as ultramer duplex oligos ordered from IDT. *dlg-1* fragments were separately amplified from pML902. The fragments were fused using overlap extension PCR to generate the final constructs. All PCR reactions were column purified (High Pure PCR Product Purification kit, cat#11732668001, Sigma Aldrich). 2 ng/μl of purified PCR products were injected into either N2 or SM507 strains along with 100 ng/μl of pRF4 plasmid as a selection marker to create transgenic lines with extrachromosomal arrays.

### smFISH, immunostaining, and microscopy

smFISH was adapted from (Tsanov et al., 2016). Custom Stellaris smFISH probes labeled with Quasar 570 dye were designed against *par-3, par-6, hmp-1*, and *erm-1* mRNAs using the Stellaris FISH Probe Designer (Biosearch Technologies, Petaluma, CA). Probes against other mRNAs were designed following the smiFISH approach as previously described (Tsanov et al., 2016). Each open reading frame was run through the Oligostan script in RStudio and 12 - 24 IDT primary smFISH probes were ordered for each mRNA (100 µM in IDTE, pH 8.0; IDT). For a full list of primary probe sequences, see Table S3. Secondary probes (FLAP-Y) with a 5’-acrydite modification and a 3’-Atto565 or a 3’-Atto637 labels were ordered from IDT. An equimolar amount of each set of primary probes was pooled in a 1.5 ml Eppendorf tube and diluted 5 times with IDTE (pH 8.0) to reach a final concentration of 0.833 µM per probe. An *in vitro* pre-hybridization reaction was set up as follow: 4 µl of primary probe-set pool, 1 µl of secondary FLAP-Y probe, 2 µl of 10x NEBuffer 3 (cat#B7003, New England Biolabs), and 13 µl of water were incubated in a thermocycler (cat#EP950040025, Eppendorf) at 85°C for 3 minutes, 65°C for 3 minutes, and 25°C for 5 minutes. Pre-hybridized FLAP-Y smFISH probes could be placed at 4°C for storage. One 6 cm plate with gravid adults and laid embryos with little bacteria lawn left were washed with 1 ml of water. Adults and larvae were discarded. An additional 1 ml of water was added to the plate. Laid embryos were gently scrubbed off with a gloved finger and transferred to a 1.5 ml Eppendorf tube. A gentle “short” 6 second spin was applied to the Eppendorf tube to pellet the embryos to minimize stress. Extra liquid was removed and embryos were allowed to rest for 10 minutes. Embryos were transferred on poly-L-lysine-coated (cat#P8920, Sigma) slides (cat#ER-303B-CE24, Thermo Scientific) and allowed to settle. Excess water was removed and 50 µl fix (1% PFA in PBS with 0.05% Triton) was added and incubated for 15 minutes. After removing the fixative solution and adding a coverslip, slides were quickly transferred on a metal plate on dry ice and stored at -80°C over-night. After a freeze-crack, slides were immediately transferred in a Coplin jar with ice-cold methanol for 5 minutes. Subsequent washes with PBS (5 minutes), PBS with 0.5% Tween-20 (10 minutes, and 20 minutes), and again PBS (5 minutes) were applied to the slides. 100 µl of hybridization solution (dextran sulfate (10% W/V) in 1 part of Formamide, 1 of 20x SSC, and 8 of water) were applied to the sample area and slides were then transferred in a humidity chamber and incubated for 1 hour at 37°C. After removal of the hybridization solution, 50 µl of new hybridization solution containing 1 µl of pre-hybridized FLAP-Y smFISH probes or 0.5 µl of Stellaris probes were applied to the sample area and slides were incubated again in a humidity chamber at 37°C for 4 hours in the dark. After incubation, hybridization solution was wicked off, and samples were washed twice with wash buffer (1 part of formamide, 1 of 20x SSC, and 8 of water). Slides with 100 µl of wash buffer were incubated for 1 hour at 37°C in a humidity chamber in the dark. The wash buffer was finally wicked off and samples were washed twice with wash buffer and mounted with 12 µl VECTASHIELD® Antifade Mounting Medium with DAPI (H-1200, Vector Laboratories). For AJM-1 antibody staining (MH27, DSHB, 1:100; Francis and Waterston, 1991) coupled to smFISH was needed, primary antibodies were added to the hybridization solution during the 4-hour incubation, and secondary antibodies (Alexafluor 546 goat anti-mouse: A-11030, Invitrogen, 1:250) to the wash buffer in the last 1-hour incubation. A widefield microscope FEI “MORE” with total internal reflection fluorescence (TIRF) and a Hamamatsu ORCA flash 4.0 cooled sCMOS camera and a Live Acquisition 2.5 software was used for capturing images. Pictures were deconvolved with the Huygens software and then processed in OMERO (https://www.openmicroscopy.org/omero/) or ImageJ (https://imagej.net/). Figures were prepared in Adobe Illustrator (https://www.adobe.com/).

### Image analysis and quantification

ImageJ (https://imagej.net/) was used for image analysis and quantitation. A script was developed to quantify *dlg-1* and *ajm-1* mRNA (Fig. 2B). The data were obtained by marking seam cell boundaries (DLG-1-GFP in the green channel) and counting mRNA dots in the red (*ajm-1* mRNA) and in the far-red (*dlg-1* mRNA) channels within the cell (cytoplasmic mRNA) and at the cell boundary (membrane localized mRNA). Transversal projections were obtained by marking the circumference of the lateral membrane of seam cells, and applying the “reslice” function (Fig. S2B’-C’). Intensity profile analyses were conducted drawing a segmented line along the middle (apicobasal profiles) or at the apical side (apical profiles) of the images shown in Fig. 5D-I. Line width was chosen accordingly to cell sizes. The “Multi Plot” function was used to generate data for the fluorescent intensities from the different channels and then processed in excel. mRNA fluorescent intensities located in the area corresponding to the nucleus were elicited from the analyses as representing transcription sites.

### Sequence analysis

To identify putative signal peptide sequences in amino acid sequences of selected proteins, we took advantage of the SignalIP-5.0 Server (Center of Biological Sequence nalysis – CBS). Nucleotide or amino acid sequences analyses were performed with Expasy (Swiss Institute of Bioinformatic – SIB) or Clustal Omega (European Molecular Biology Laborytory-European Bioinformatic Institute – EMBL-EBI).

## Acknowledgments

We thank Jeff Hardin and his lab for sharing their *dlg-1*::GFP transgenic lines, Laurent Guerard and the Imaging Core Facility of the Biozentrum for support in imaging analyses, Poulomi Ray for new transgenic *dlg-1*::GFP lines and inputs, and the whole Mango lab for scientific discussions. Some strains were provided by the CGC, which is funded by NIH Office of Research Infrastructure Programs (P40 OD010440).

## Supplemental material

### Supplemental table legends

**Table S1. List of methionine amino acid within the DLG-1 sequence**. Amino acid positions and domain location of the 11 methionine amino acids found in the DLG-1 sequence, corresponding to the green asterisks in Fig. 4A.

**Table S2. List of *C. elegans* strains**. Detailed list of names, genotypes, and references of *C. elegans* strains used in this work.

**Table S3. List of smFISH probes**. Sequences of the smFISH primary probes used in this study.

### Supplemental figure legends

**Supplemental figure 1.**
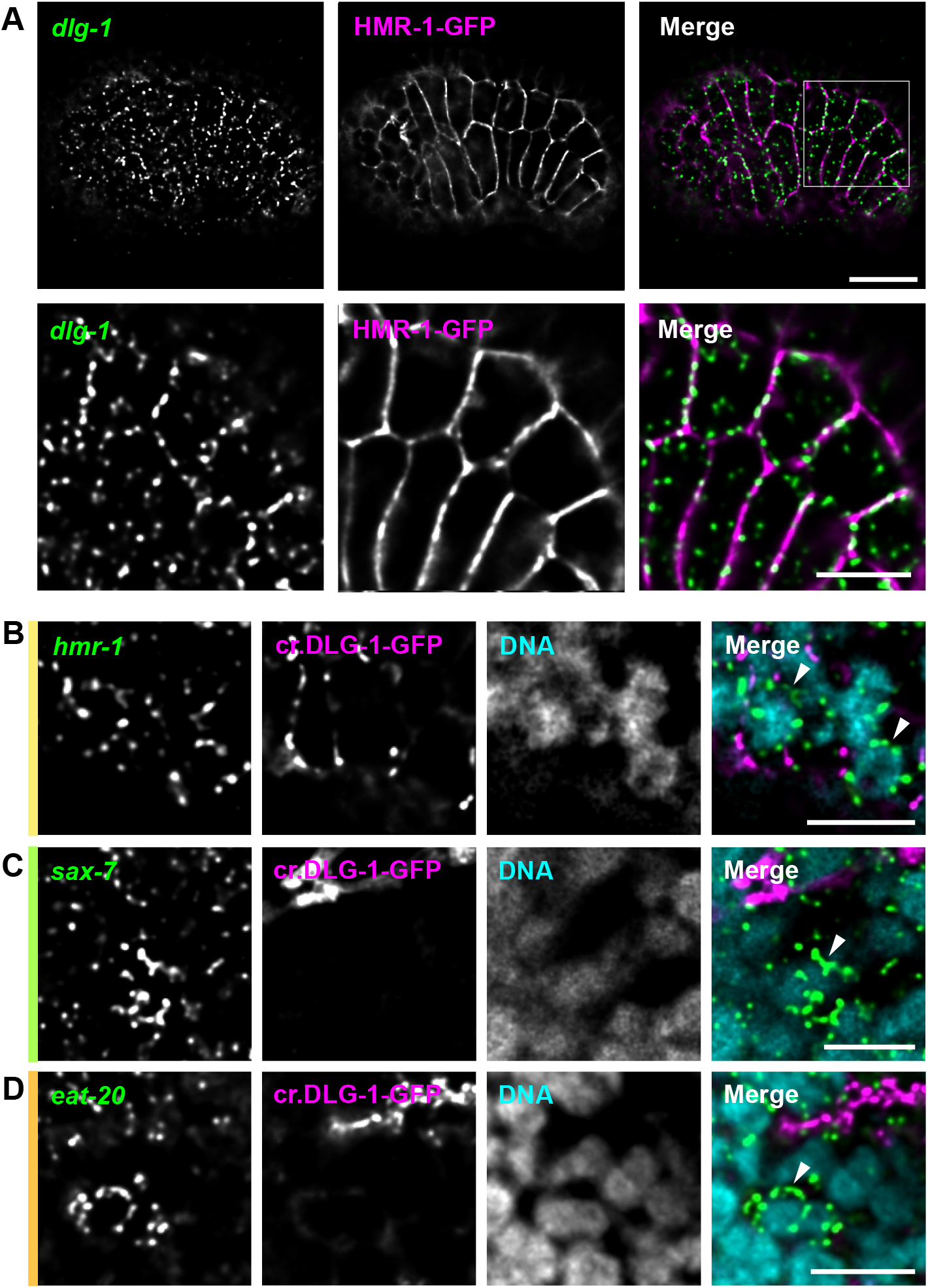
Endogenous *dlg-1* is enriched at the *Ce*AJ; transmembrane protein-coding mRNAs show instances of perinuclear localization. **A**. Fluorescent micrographs of the entire *C. elegans* embryos (upper panels) and zoom-ins (lower panels) showing smFISH signal of endogenous *dlg-1* mRNAs (green) in epidermal and seam cells of a bean stage, fluorescent signal of the transgenic GFP-tagged HMR-1 protein (HMR-1-GFP, magenta), and merges. Scale bar (upper panels): 10 µm. Scale bars (lower panels): 5 µm. **B-D**. Fluorescent micrographs of portions of *C. elegans* embryos showing instances of perinuclearly localized mRNAs coding for transmembrane proteins (HMR-1, SAX-7, and EAT-20). smFISH signal of localized mRNAs *hmr-1* (B), *sax-7* (C), and *eat-20* (D) (green), fluorescent signal of the CRISPR-engineered DLG-1-GFP (cr.DLG-1-GFP) (magenta), and merges. On the left side of each image, bars with the same color-code as in Fig. 1A to indicate to which sub-class of factors the corresponding mRNA codes for. Scale bars: 5 µm.

**Supplemental figure 2.**
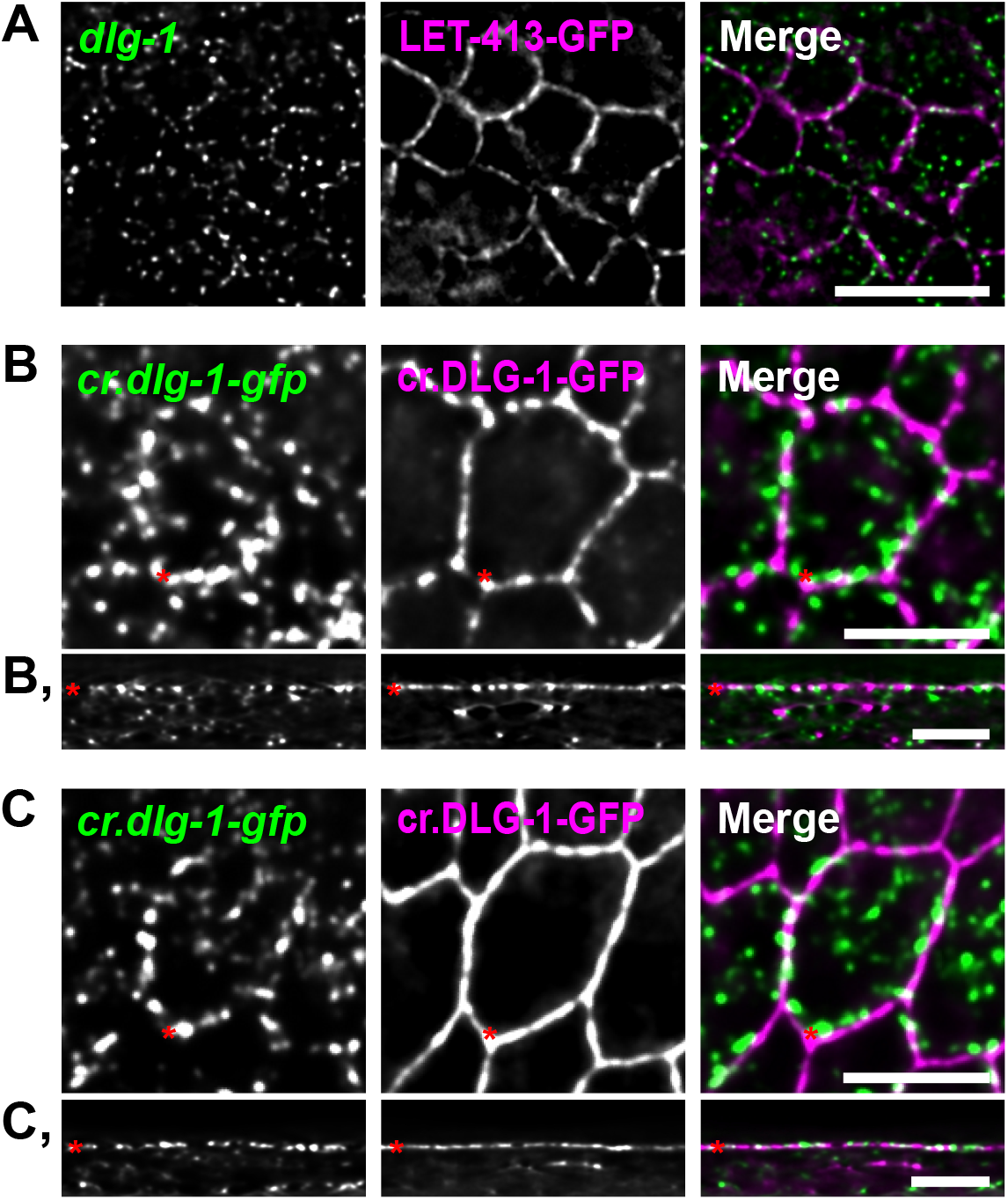
*dlg-1* mRNA is enriched at the cell membrane prior to *Ce*AJ maturation. **A**. Fluorescent micrographs of developing epithelial cells of a portion of a *C. elegans* embryo at the 4E stage showing smFISH signal of the endogenous *dlg-1* mRNA (green), fluorescent signal of the CRISPR-engineered GFP-tagged DLG-1 protein transgenic GFP-tagged LET-413 protein (LET-413-GFP, magenta), and merges. Scale bars: 10 µm. **B**. Top view of fluorescent micrographs of a seam cell at the embryonic bean stage. Images show smFISH signal of the *dlg-1* mRNA (*cr*.*dlg-1-gfp*, green), fluorescent signal of the CRISPR-engineered GFP-tagged DLG-1 protein (cr.DLG-1-GFP, magenta), and merges. Red asterisks mark the site from which transverse projections shown in (B’) start from. Scale bar: 5 µm. **B’**. Corresponding transverse sections on a flat plane of the sole rolled out membrane of the seam cell shown in (B). Red asterisks mark the horizontal location of *Ce*AJ and the starting site shown in (B). Scale bar: 5 µm. **C-C’**. Same as in (B-B’), but showing a seam cell at the embryonic comma stage.

**Supplemental figure 3.**
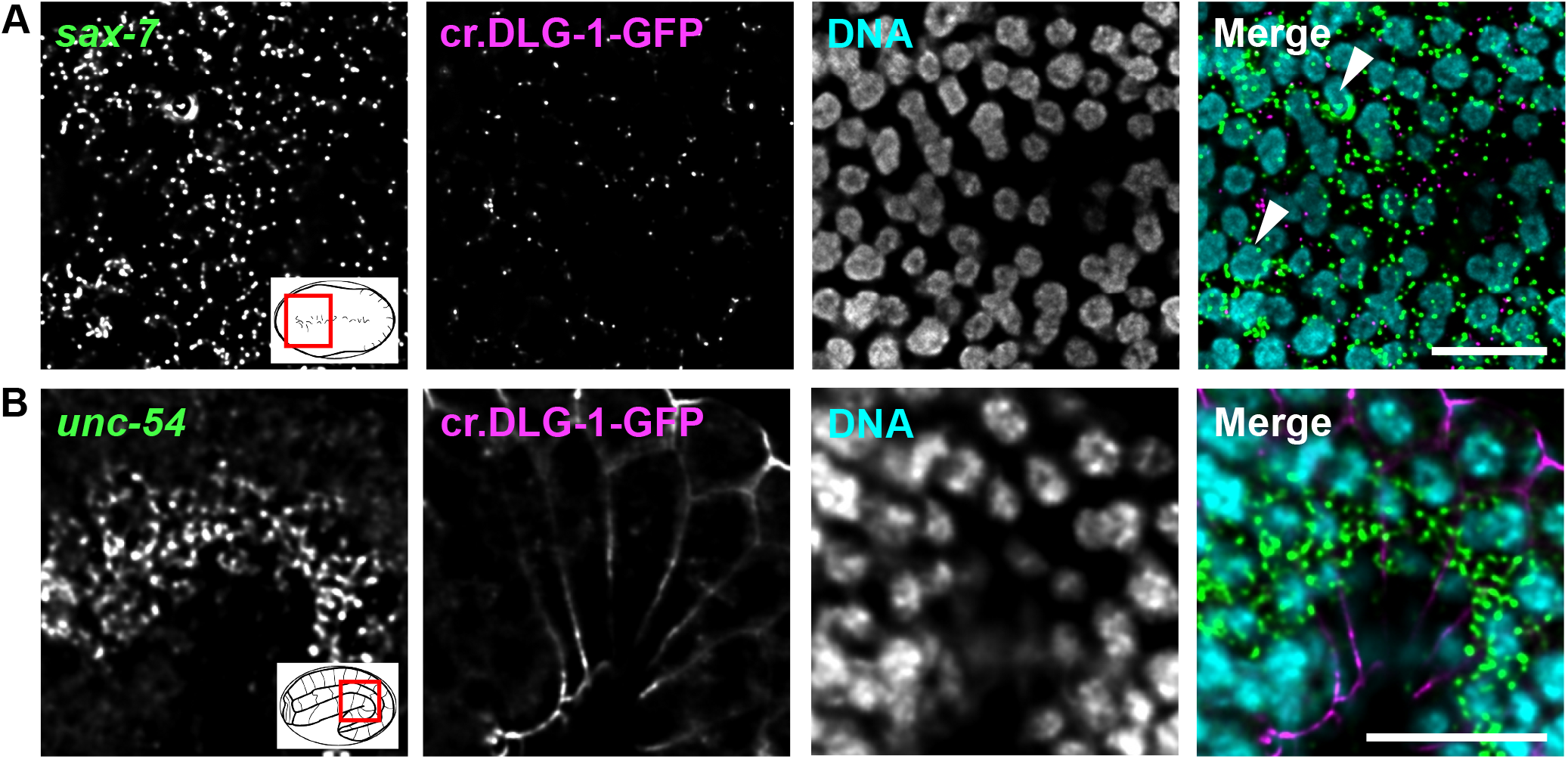
The UTRs enrolled in transgenic lines derive from unlocalized mRNAs. **A-B**. Maximum intensity projections of 5 (1.08 µm) (A) and 3 (0.54 µm) (B) Z-stacks of fluorescent micrographs of portions (red box in the cartoon) of *C. elegans* embryos at the 16E (A) and 1.5-fold (B) stages showing smFISH signal of *sax-7* (A) and *unc-54* (B) mRNAs (green), fluorescent signal of the endogenous GFP-tagged DLG-1 protein (cr.DLG-1-GFP, magenta), DNA (cyan), and merges. Scale bars: 10 µm.

**Supplemental figure 4.**
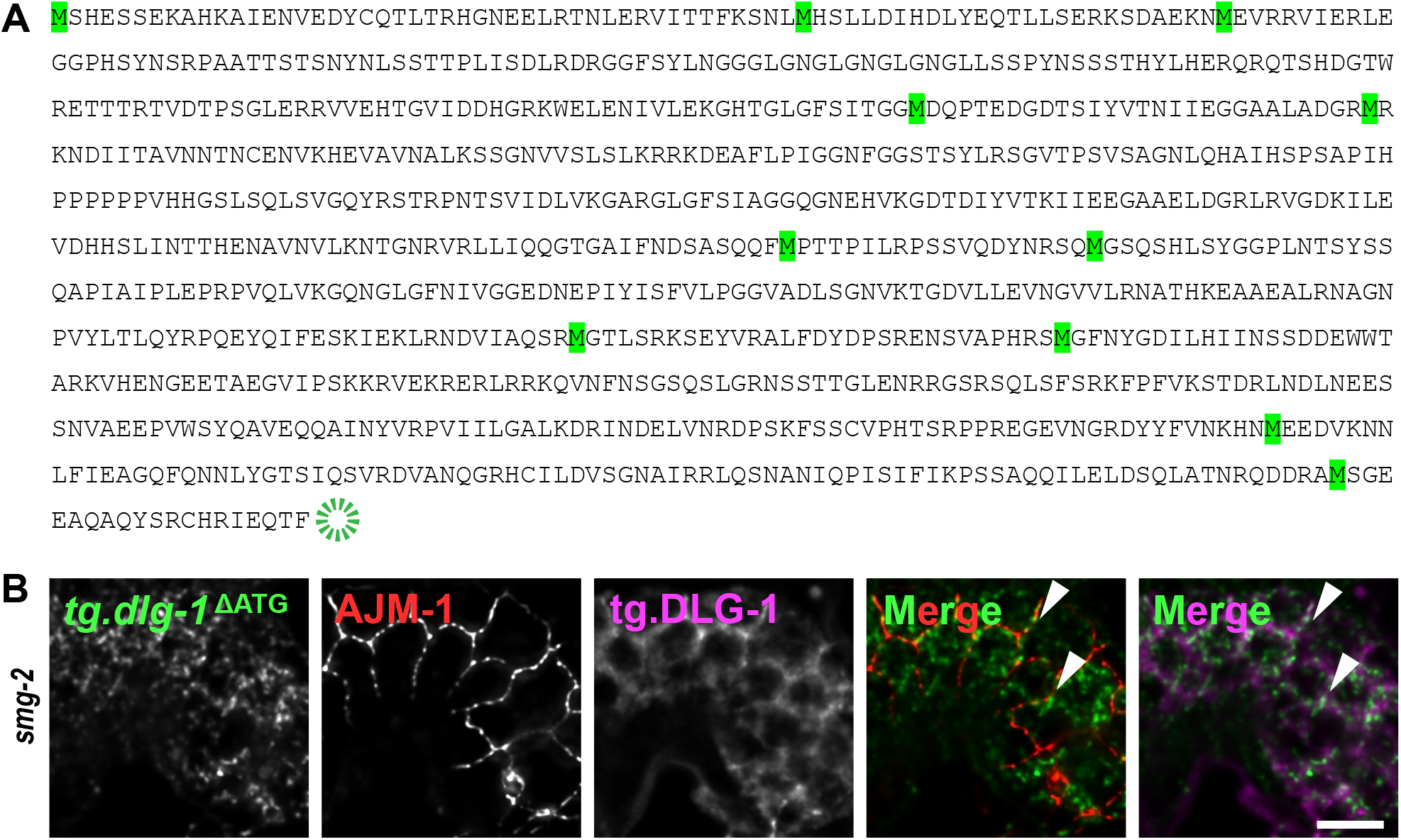
*dlg-1* mRNA coding sequence possesses putative alternative start codons that may allow its partial translation. **A**. Transgenic DLG-1 protein sequence. Highlighted in green are methionine amino acids. **B**. Fluorescent micrographs of a lateral portion of seam and epithelial cells at the late bean stage of a *C. elegans* embryo showing an example of a “ΔATG” transgene that is able to express its coded protein. As in Fig. 4: smFISH signal of altered ATG (“ΔATG”) *tg*.*dlg-1-gfp* mRNAs (green), immuno-fluorescent signal of the endogenous AJM-1 protein (red), fluorescent signal of the corresponding ΔATG tg.DLG-1-GFP protein (magenta), and merges. Arrowheads: examples of laterally localized mRNAs. Scale bar: 5 µm.

**Supplemental figure 5.**
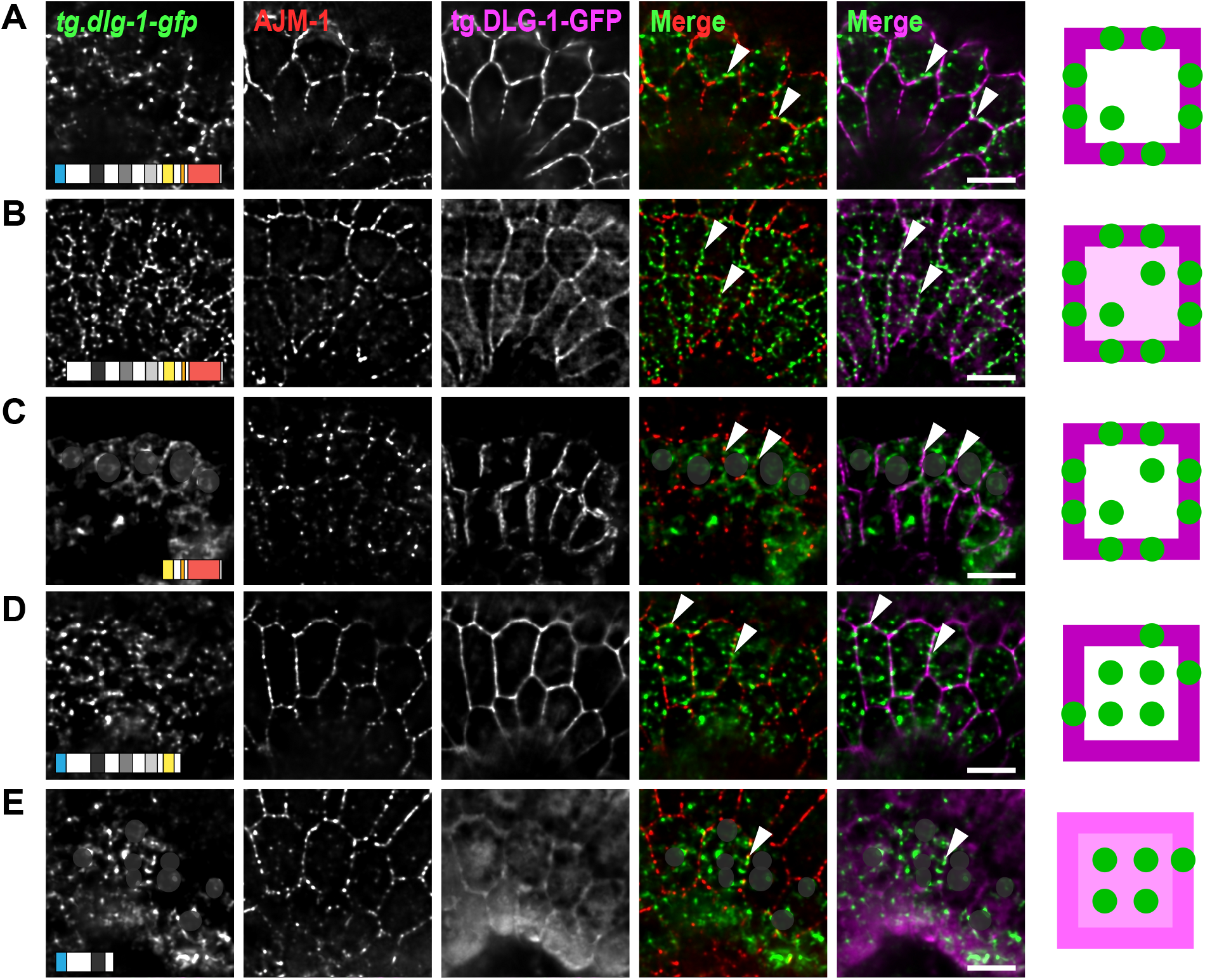
Specific domain-coding sequence of *dlg-1* mRNA are required for its normal lateral localization. **A-E**. Top views of fluorescent micrographs of a lateral portion of seam and epidermal cells at the late bean stage of *C. elegans* embryos showing smFISH signal of transgenic *tg*.*dlg-1-gfp* mRNAs (full length (A), ΔL27 (B), SH3-cHk-GuK (C), ΔcHk-GuK (D), and L27-PDZ1 (E); (green)), immuno-fluorescent signal of the endogenous AJM-1 protein (red), fluorescent signal of the transgenic GFP-tagged DLG-1 protein coded by the corresponding transgene (magenta), and merges. Shaded gray circles locate the nuclear regions. Arrowheads: examples of laterally localized mRNAs. Scale bars: 5 µm. Right side: simplified schematics based on the fluorescent images on the right. Top view of a seam cell (square) modelling transgenic mRNA and protein localizations. Green circles: transgenic *dlg-1* mRNA. Shades of magenta: varying degrees of transgenic DLG-1 protein (lighter magenta – lower amount of protein; darker magenta – higher amount of protein) along the membrane (borders) and in the cytoplasm (middle part).

## References

Armenti, S. T., & Nance, J. (2012). Adherens junctions in C. elegans embryonic morphogenesis. Subcell Biochem, 60, 279-299. https://doi.org/10.1007/978-94-007-4186-7_12

Bernadskaya, Y. Y., Patel, F. B., Hsu, H. T., & Soto, M. C. (2011). Arp2/3 promotes junction formation and maintenance in the Caenorhabditis elegans intestine by regulating membrane association of apical proteins. Mol Biol Cell, 22(16), 2886-2899. https://doi.org/10.1091/mbc.E10-10-0862

Bossinger, O., Klebes, A., Segbert, C., Theres, C., & Knust, E. (2001). Zonula adherens formation in Caenorhabditis elegans requires dlg-1, the homologue of the Drosophila gene discs large. Dev Biol, 230(1), 29–42. https://doi.org/10.1006/dbio.2000.0113

Bossinger, O., Wiesenfahrt, T., & Hoffmann, M. (2015). Establishment and Maintenance of Cell Polarity in the C. elegans Intestine. In: Ebnet K. (eds) Cell Polarity 2. Springer, Cham. https://doi.org/10.1007/978-3-319-14466-5_2

Brenner, S. (1974). The genetics of Caenorhabditis elegans. Genetics, 77(1), 71–94.

Broadus, J., Fuerstenberg, S., & Doe, C. Q. (1998). Staufen-dependent localization of prospero mRNAcontributes to neuroblast daughter-cell fate. Nature, 391, 792–795.

Chaudhuri, A., Das, S., & Das, B. (2020). Localization elements and zip codes in the intracellular transport and localization of messenger RNAs in Saccharomyces cerevisiae. Wiley Interdiscip Rev RNA, 11(4), e1591. https://doi.org/10.1002/wrna.1591

Chen, L., & Zhou, S. (2010). “CRASH”ing with the worm: insights into L1CAM functions and mechanisms. Dev Dyn, 239(5), 1490–1501. https://doi.org/10.1002/dvdy.22269

Chouaib, R., Safieddine, A., Pichon, X., Imbert, A., Kwon, O. S., Samacoits, A., Traboulsi, A. M., Robert, M. C., Tsanov, N., Coleno, E., Poser, I., Zimmer, C., Hyman, A., Le Hir, H., Zibara, K., Peter, M., Mueller, F., Walter, T., & Bertrand, E. (2020). A Dual Protein-mRNA Localization Screen Reveals Compartmentalized Translation and Widespread Co-translational RNA Targeting. Dev Cell, 54(6), 773–791 e775. https://doi.org/10.1016/j.devcel.2020.07.010

Costa, M., Raich, W., Agbunag, C., Leung, B., Hardin, J., & Priess, J. R. (1998). A putative catenin-cadherin system mediates morphogenesis of the Caenorhabditis elegans embryo. J Cell Biol, 141, 297–308.

Ephrussi, A., Dickinson, L. K., & Lehmann, R. (1991). oskar Organizes the Germ Plasm and Directs Localization of the Posterior Determinant nanos. Cell, 66, 37–50.

Firestein, B. L., & Rongo, C. (2001). DLG-1 Is a MAGUK Similar to SAP97 and Is Required for Adherens Junction Formation. Mol Biol Cell, 12, 3465–3475.

Frigerio, G., Burri, M., Bopp, D., Baumgartner, S., & Noll, M. (1986). Structure of the segmentation gene paired and the Drosophila PRD gene set as part of a gene network. Cell, 47, 735–746.

Gallo, C. M., Munro, E., Rasoloson, D., Merritt, C., & Seydoux, G. (2008). Processing bodies and germ granules are distinct RNA granules that interact in C. elegans embryos. Dev Biol, 323(1), 76–87. https://doi.org/10.1016/j.ydbio.2008.07.008

Ghosh, S., Marchand, V., Gaspar, I., & Ephrussi, A. (2021). Control of RNP motility and localization by splicing-dependent structure in oskar mRNA. Nat Struct Mol Biol, 19(4), 441–9. https://doi:10.1038/nsmb.2257.

Gobel, V., Barrett p. L., Hall D. H., & Fleming J. T. (2004). Lumen morphogenesis in C. elegans requires the membrane-cytoskeleton linker erm-1. Dev Cell, 6(6), 865–873. https://doi.org/10.1016/j.devcel.2004.05.018

Hachet, O., & Ephrussi, A. (2004). Splicing of oskarRNA in the nucleus is coupled to its cytoplasmic localization. Nature, 428(6986), 959–963. https://doi.org/10.1038/nature02484

Heppert, J. K., Pani, A. M., Roberts, A. M., Dickinson, D. J., & Goldstein, B. (2018). A CRISPR Tagging-Based Screen Reveals Localized Players in Wnt-Directed Asymmetric Cell Division. Genetics, 208(3), 1147–1164. https://doi.org/10.1534/genetics.117.300487

Hermesh, O., & Jansen, R. p. (2013). Take the (RN)A-train: localization of mRNA to the endoplasmic reticulum. Biochim Biophys Acta, 1833(11), 2519–2525. https://doi.org/10.1016/j.bbamcr.2013.01.013

Hirashima, T., Tanaka, R., Yamaguchi, M., & Yoshida, H. (2018). The ABD on the nascent polypeptide and PH domain are required for the precise Anillin localization in Drosophila syncytial blastoderm. Sci Rep, 8(1), 12910. https://doi.org/10.1038/s41598-018-31106-0

Jambhekar, A., & Derisi, J. L. (2007). Cis-acting determinants of asymmetric, cytoplasmic RNA transport. RNA, 13(5), 625–642. https://doi.org/10.1261/rna.262607

Jambor, H., Surendranath, V., Kalinka, A. T., Mejstrik, P., Saalfeld, S., & Tomancak, p. (2015). Systematic imaging reveals features and changing localization of mRNAs in Drosophila development. Elife, 4. https://doi.org/10.7554/eLife.05003

Katz, Z. B., Wells, A. L., Park, H. Y., Wu, B., Shenoy, S. M., & Singer, R. H. (2012). beta-Actin mRNA compartmentalization enhances focal adhesion stability and directs cell migration. Genes Dev, 26(17), 1885–1890. https://doi.org/10.1101/gad.190413.112

Kislauskis, E. H., Zhu, X., & Singer, R. H. (1994). Sequences Responsible for Intracellular Localization of β-Actin Messenger RNA Also Affect Cell Phenotype. J Cell Biol, 127, 441–451.

Kourtidis, A., Necela, B., Lin, W. H., Lu, R., Feathers, R. W., Asmann, Y. W., Thompson, E. A., & Anastasiadis, p. Z. (2017). Cadherin complexes recruit mRNAs and RISC to regulate epithelial cell signaling. J Cell Biol, 216(10), 3073–3085. https://doi.org/10.1083/jcb.201612125

Kwon, O. S., Mishra, R., Safieddine, A., Coleno, E., Alasseur, Q., Faucourt, M., Barbosa, I., Bertrand, E., Spassky, N., & Le Hir, H. (2021). Exon junction complex dependent mRNA localization is linked to centrosome organization during ciliogenesis. Nat Commun, 12(1), 1351. https://doi.org/10.1038/s41467-021-21590-w

Lecuyer, E., Yoshida, H., Parthasarathy, N., Alm, C., Babak, T., Cerovina, T., Hughes, T. R., Tomancak, P., & Krause, H. M. (2007). Global analysis of mRNA localization reveals a prominent role in organizing cellular architecture and function. Cell, 131(1), 174–187. https://doi.org/10.1016/j.cell.2007.08.003

Li, P., Yang, X., Wasser, M., Cai, Y., & Chia, W. (1997). Inscuteable and Staufen Mediate Asymmetric Localization and Segregation of prospero RNA during Drosophila Neuroblast Cell Divisions. Cell, 90, 437–447.

Lockwood, C. A., Lynch, A. M., & Hardin, J. (2008). Dynamic analysis identifies novel roles for DLG-1 subdomains in AJM-1 recruitment and LET-413-dependent apical focusing. J Cell Sci, 121(Pt 9), 1477–1487. https://doi.org/10.1242/jcs.017137

McKeown, C., Praitis, V., & Austin, J. (1998). sma-1 encodes a betaH-spectrin homolog required for Caenorhabditis elegans morphogenesis. Development, 125(11), 2087–2098. https://doi.org/10.1242/dev.125.11.2087

McMahon, L., Legouis, R., Vonesch, J. L., & Labouesse, M. (2001). Assembly of C. elegans apical junctions involves positioning and compaction by LET-413 and protein aggregation by the MAGUK protein DLG-1. J Cell Sci, 114, 2265–2277.

Moor, A. E. G. M.,; Massasa, E. E.; Lemze, D.; Weizman, T.; Shenhav, R.; Baydatch, S.; Mizrahi, O.; Winkler, R.; Golani, O.; Stern-Ginossar, N.; Itzkovitz, S. (2017). Global mRNA polarization regulates translation efficiency in the intestinal epithelium. Science, 357, 1299–1303.

Nagaoka, K., Udagawa, T., & Richter, J. D. (2012). CPEB-mediated ZO-1 mRNA localization is required for epithelial tight-junction assembly and cell polarity. Nat Commun, 3, 675. https://doi.org/10.1038/ncomms1678

Nyathi, Y., Wilkinson, B. M., & Pool, M. R. (2013). Co-translational targeting and translocation of proteins to the endoplasmic reticulum. Biochim Biophys Acta, 1833(11), 2392–2402. https://doi.org/10.1016/j.bbamcr.2013.02.021

Ouyang, J. p.T., Folkmann, A., Bernard, L., Lee C. Y., Seroussi, U., Charlesworth A. G., Claycomb J. M., & Seydoux, G. (2019). P Granules Protect RNA Interference Genes from Silencing by piRNAs. Dev Cell, 50(6), 716–728 e716. https://doi.org/10.1016/j.devcel.2019.07.026

Parker, D. M., Winkenbach, L. P., Boyson, S., Saxton, M. N., Daidone, C., Al-Mazaydeh, Z. A., Nishimura, M. T., Mueller, F., & Osborne Nishimura, E. (2020). mRNA localization is linked to translation regulation in the Caenorhabditis elegans germ lineage. Development, 147(13). https://doi.org/10.1242/dev.186817

Pettitt, J., Cox, E. A., Broadbent, I. D., Flett, A., & Hardin, J. (2003). The Caenorhabditis elegans p120 catenin homologue, JAC-1, modulates cadherin-catenin function during epidermal morphogenesis. J Cell Biol, 162(1), 15–22. https://doi.org/10.1083/jcb.200212136

Rebagliati, M. R., Weeks, D. L., Harvey, R. P., & Melton, D. A. (1985). Identification and Cloning of Localized Maternal RNAs from Xenopus Eggs. Cell, 42, 769–777.

Ryder, p. V., & Lerit, D. A. (2018). RNA localization regulates diverse and dynamic cellular processes. Traffic, 19(7), 496–502. https://doi.org/10.1111/tra.12571

Safieddine, A., Coleno, E., Salloum, S., Imbert, A., Traboulsi, A. M., Kwon, O. S., Lionneton, F., Georget, V., Robert, M. C., Gostan, T., Lecellier, C. H., Chouaib, R., Pichon, X., Le Hir, H., Zibara, K., Mueller, F., Walter, T., Peter, M., & Bertrand, E. (2021). A choreography of centrosomal mRNAs reveals a conserved localization mechanism involving active polysome transport. Nat Commun, 12(1), 1352. https://doi.org/10.1038/s41467-021-21585-7

Scheckel, C., Gaidatzis, D., Wright, J. E., & Ciosk, R. (2012). Genome-wide analysis of GLD-1-mediated mRNA regulation suggests a role in mRNA storage. PLoS Genet, 8(5), e1002742. https://doi.org/10.1371/journal.pgen.1002742

Shibata, Y., Fujii, T., Dent, J. A., Fujisawa, H., & Takagi, S. (1999). EAT-20, a Novel Transmembrane Protein With EGF Motifs, Is Required for Efficient Feeding in Caenorhabditis elegans. Genetics, 154, 635–646.

Shukla, A., Yan, J., Pagano, D. J., Dodson, A. E., Fei, Y., Gorham, J., Seidman, J. G., Wickens, M., & Kennedy, S. (2020). poly(UG)-tailed RNAs in genome protection and epigenetic inheritance. Nature, 582(7811), 283–288. https://doi.org/10.1038/s41586-020-2323-8

Standart, N., & Weil, D. (2018). P-Bodies: Cytosolic Droplets for Coordinated mRNA Storage. Trends Genet, 34(8), 612–626. https://doi.org/10.1016/j.tig.2018.05.005

Sulston, J. E., Schierenberg, E., White, J. G., & Thomson, J. N. (1983). The Embryonic Cell Lineage of the Nematode Caenorhabditis elegans. Dev Biol, 100, 64–119.

Takizawa, p. A. S. A.; Swedlow, J. R.;, & Herskowitz, I. V. R. D. (1997). Actin-dependent localization of an RNA encoding a cell-fate determinant in yeast. Nature, 389, 90–93.

Tsanov, N., Samacoits, A., Chouaib, R., Traboulsi, A. M., Gostan, T., Weber, C., Zimmer, C., Zibara, K., Walter, T., Peter, M., Bertrand, E., & Mueller, F. (2016). smiFISH and FISH-quant - a flexible single RNA detection approach with super-resolution capability. Nucleic Acids Res, 44(22), e165. https://doi.org/10.1093/nar/gkw784

Van Furden, D., Johnson, K., Segbert, C., & Bossinger, O. (2004). The C. elegans ezrin-radixin-moesin protein ERM-1 is necessary for apical junction remodelling and tubulogenesis in the intestine. Dev Biol, 272(1), 262–276. https://doi.org/10.1016/j.ydbio.2004.05.012

Voronina, E. (2013). The diverse functions of germline P-granules in Caenorhabditis elegans. Mol Reprod Dev, 80(8), 624–631. https://doi.org/10.1002/mrd.22136

Weis, B. L., Schleiff, E., & Zerges, W. (2013). Protein targeting to subcellular organelles via MRNA localization. Biochim Biophys Acta, 1833(2), 260–273. https://doi.org/10.1016/j.bbamcr.2012.04.004

Wright, J. E., Gaidatzis, D., Senften, M., Farley, B. M., Westhof, E., Ryder, S. P., & Ciosk, R. (2011). A quantitative RNA code for mRNA target selection by the germline fate determinant GLD-1. EMBO J, 30(3), 533–545. https://doi.org/10.1038/emboj.2010.334

Yan, D., Wu, Z., Chisholm, A. D., & Jin, Y. (2009). The DLK-1 kinase promotes mRNA stability and local translation in C. elegans synapses and axon regeneration. Cell, 138(5), 1005–1018. https://doi.org/10.1016/j.cell.2009.06.023

Yergert, K. M., Doll, C. A., O’Rouke, R., Hines, J. H., & Appel, B. (2021). Identification of 3’ UTR motifs required for mRNA localization to myelin sheaths in vivo. PLoS Biol, 19(1), e3001053. https://doi.org/10.1371/journal.pbio.3001053

## Supplemental references

Marston, D. J., Higgins, C. D., Peters, K. A., Cupp, T. D., Dickinson, D. J., Pani, A. M., Moore, R. P., Cox, A. H., Kiehart, D. P., & Goldstein, B. (2016). MRCK-1 Drives Apical Constriction in C. elegans by Linking Developmental Patterning to Force Generation. Curr Biol, 26(16), 2079–89. https://doi:10.1016/j.cub.2016.06.010.

Jia, R., Li, D., Li., M., Chai, Y., Liu, Y., Xie, Z., Shao, W., Xie, C., Li, L., Huang, X., Chen, L., Li, W., & Ou, G. (2019). Spectrin-based membrane skeleton supports ciliogenesis. PLoS Biol, 17(7), e3000369. https://doi:10.1371/journal.pbio.3000369.

Quintin, S., Wang, S., Pontabry, J., Bender, A., Robin, F., Hyenne, V., Landmann, F., Gally, C., Oegema, K., & Labouesse, M. (2016). Non-centrosomal epidermal microtubules act in parallel to LET-502/ROCK to promote C. elegans elongation. Development, 143(1), 160–73. https://doi.10.1242/dev.126615.

Yang, Y., Zhang, Y., Li, W. J., Jiang, Y., Zhu, Z., Hu, H., Li, W., Wu, J. W., Wang, Z. X., Dong, M. Q., Huang, S., & Ou, G. (2017). Spectraplakin Induces Positive Feedback between Fusogens and the Actin Cytoskeleton to Promote Cell-Cell Fusion. Dev Cell, 41(1), 107–120. https://doi:10.1016/j.devcel.2017.03.006.

Achilleos, A., Wehman, A. M., & Nance, J. (2010). PAR-3 mediates the initial clustering and apical localization of junction and polarity proteins during C. elegans intestinal epithelial cell polarization. Development, 137(11), 1833–42. https://doi:10.1242/dev.047647.

Page, M. F., Carr, B., Anders, K. R., Grimson, A., & Anderson, p. (1999) SMG-2 is a phosphorylated protein required for mRNA surveillance in Caenorhabditis elegans and related to Upf1p of yeast. Mol Cell Biol, 19(9), 5943–51. https://doi:10.1128/mcb.19.9.5943.

